# Macrophage EHD1 promotes inflammation and stabilizes sortilin to accelerate atherosclerosis

**DOI:** 10.1101/2025.10.27.684958

**Authors:** Fanglin Ma, Yu Liu, Yang Xu, Wenli Liu, Neha Gupta, Yiwei Zhu, Lieying Xiao, James G. Traylor, Oren Rom, Jason C. Kovacic, Trevor P. Fidler, Arif Yurdagul, A. Wayne Orr, Xin Huang, Bishuang Cai

## Abstract

**Background:** Macrophages are key players in the pathogenesis of atherosclerosis. They trigger immune responses through their cell-surface receptors. However, how macrophages regulate those receptors in response to pro-inflammatory stimuli is not completely understood. Endocytic membrane trafficking involving receptor internalization, followed by endosomal transport and recycling of the internalized receptors, plays essential roles in balancing cell-surface receptors to meet cellular needs. Here, we explored the role of the endocytic regulator EHD1 in immune responses in macrophages and determined its contribution to atherosclerosis progression.

**Methods:** EHD1 expression profiles in mouse and human plaques were determined by single-cell RNA sequencing (scRNA-seq) and immunofluorescence staining. Bone marrow transplantation (BMT) by transplanting bone marrow cells from *Ehd1^−/−^* or littermate wild-type mice to irradiated *Ldlr^−/−^* mice was performed to determine the effect of EHD1 deletion on atherosclerosis progression. *In vitro* mechanistic studies including inflammation signaling and endocytosis assays were performed in bone marrow–derived macrophages.

**Results:** EHD1 expression in macrophages is enhanced as atherosclerosis progresses in both mice and humans. Histological analysis of aortic root sections from BMT mice showed that EHD1 deletion reduces lesion size. ScRNA-seq of aortic CD45^+^ cells demonstrated that EHD1 deletion attenuates pro-inflammatory responses and cell–cell interactions. Mechanistic studies revealed that EHD1 accelerates the endocytic recycling of TNFR2 and activates NF-kB, leading to increased expression of inflammatory cytokines. Moreover, EHD1 interacts with retromer and stabilizes sortilin, a retrograde cargo of retromer and a risk factor for atherosclerosis.

**Conclusions:** EHD1 promotes inflammation by enhancing TNFR2−NF-kB signaling and stabilizing sortilin, leading to accelerated atherosclerosis. Our study reveals novel roles for EHD1-mediated membrane trafficking in macrophage function and paves the way to innovative therapeutic strategies that aim to address dysregulated membrane trafficking in atherosclerosis.

## Introduction

Atherosclerosis is a chronic inflammatory disease characterized by the infiltration of various immune cells.^1,2^ As multifunctional cells of the innate immune system, infiltrating monocyte-derived macrophages are major contributors to the initiation, progression, and destabilization of atherosclerotic plaques.^3^ In response to cardiovascular injury, inflammatory Ly6C^high^ monocytes infiltrate the injured vessel wall and differentiate into macrophages. These macrophages trigger pro-inflammatory cascades that further amplify leukocyte recruitment, leading to plaque progression.^4^ The activation of nuclear factor-kB (NF-kB) is a crucial mechanism for initiating and sustaining a pro-inflammatory response in macrophages. As a transcription factor, NF-kB drives the transcription of genes for inflammatory cytokines, chemokines, and adhesion mediators, leading to immune cell recruitment and amplified inflammation.^5^ Clinical studies have confirmed that excessive inflammation remains a significant residual risk for atherosclerotic cardiovascular disease, despite optimal cholesterol-lowering therapies.^6^ Hence, clinical trials targeting excessive inflammation using drugs to inhibit pro-inflammatory cytokines, such as IL-1β (canakinumab) and IL-6 (ziltivekimab), have been shown to reduce cardiovascular events in patients.^7^ Given the essential role of inflammation in the pathogenesis of atherosclerosis, identifying new mechanisms that regulate inflammation in macrophages is of paramount importance.

Macrophages rely heavily on cell-surface proteins such as receptors and transporters for their immune response.^8^ Therefore, studying the turnover of cell-surface proteins in macrophages is essential. Endocytic membrane trafficking is a key cellular process that controls the levels of receptors and other plasma membrane proteins on the cell surface. Upon internalization, receptors and other transmembrane proteins are segregated into budding vesicles that are cleaved from the plasma membrane and trafficked to a key endocytic compartment known as the early endosome.^9^ From early endosomes, cargoes may be transported to late endosomes and lysosomes for degradation, or to recycling endosomes for recycling back to the plasma membrane.^9,10^ C-terminal Eps15 Homology Domain (EHD) proteins—EHD1, EHD2, EHD3, and EHD4—are evolutionarily conserved regulators in endocytic trafficking.^11^ They are localized to intracellular endosomes and regulate endosomal trafficking and protein sorting. Among these four EHD proteins, EHD1 is the best characterized. EHD1 is well-known to facilitate the endocytic recycling of transmembrane cargoes from the recycling endosome to the plasma membrane.^12–17^ In addition to recycling, EHD1 also promotes retromer-mediated retrograde transport from endosomes to the trans-Golgi network (TGN).^18,19^ EHD2 is involved in internalization,^20,21^ and EHD3 and EHD4 regulate cargo exit from early endosomes.^22–24^ Emerging evidence suggests that EHD proteins play important roles in cardiovascular diseases.^25^ For instance, EHD-mediated endocytic membrane trafficking has been shown to regulate ion channels and gap junctions in cardiomyocytes in the context of arrhythmias and myocardial infarction.^26,27^ However, the roles of EHD proteins in macrophage biology and atherosclerosis are unknown.

Here, we demonstrate that among all the EHD-family proteins, EHD1 is preferentially expressed in pro-inflammatory macrophages, and its expression is positively correlated with pro-inflammatory cytokines in mouse and human atherosclerotic plaques. Hematopoietic EHD1 deletion suppresses plaque inflammation and progression. Single-cell RNA sequencing (scRNA-seq) of aortic immune cells revealed that EHD1 deletion suppresses inflammatory response in aortic macrophages and reduces the interaction of macrophages with other immune cells. Mechanistically, we found that EHD1 promotes the endocytic recycling of tumor necrosis factor receptor 2 (TNFR2) to the cell surface, potentially leading to the activation of NF-kB. Moreover, EHD1 stabilizes sortilin, a retrograde cargo and a risk factor for cardiovascular disease via its role in inducing cytokine secretion.^28^ Together, these findings elucidate a novel role of macrophage EHD1 in accelerating inflammation during atherosclerosis.

## Methods

### Experimental animals

Male wildtype (WT) C57BL/6J (#000664, 7 weeks old), male *Ldlr*^tm1Her^/J (*Ldlr^-/-^*, #002207, 7 weeks old), and *Ehd1*^tm1Mhor^/J mice of both genders (*Ehd1^-/-^*, #023073, 7 weeks old) were purchased from Jackson Laboratory (Bar Harbor, ME). The mice were allowed to adapt to housing in the animal facility for 1 week before the experiments began. For the atherosclerosis studies, bone marrow cells from male *Ehd1^-/-^* mice and sex-matched WT littermate mice were transplanted into *Ldlr^−/−^*male mice, respectively, as previously described.^29^ Briefly, *Ldlr^−/−^* recipients were irradiated using a cesium gamma source at a dose of 1,000 rads. After 5 h rest, irradiated *Ldlr^−/−^* mice were injected with 5 x 10^6^ bone marrow cells isolated from either *Ehd1^-/-^*or WT donors. Following transplantation, the mice were maintained on antibiotic water and a normal diet for 6 weeks, after which they were switched to regular water and a Western-type diet (WD; Harlan Teklad, TD88137) for 12 weeks to induce atherosclerosis. During the WD feeding period, mouse body weight was recorded weekly. Five mice per cage were housed in standard cages at 22°C in a 12– 12 h light-dark cycle in a barrier facility. All mice were euthanized via exsanguination under isoflurane anesthesia. Blood was obtained through vena cava puncture, subsequently centrifuged at 13,000 × g for 10 min. The resulting plasma samples were collected and stored at -80°C for subsequent biochemical analysis. All procedures were performed according to protocols approved by the Animal Care and Use Committee of Icahn School of Medicine at Mount Sinai.

### Human coronary artery section staining

Human coronary arteries were isolated during routine autopsy at LSU Health Shreveport as part of the Louisiana Coronary Artery (LOCATE) Biobank. Deemed nonhuman research by the local IRB due to the use of postmortem tissue exclusively (STUDY00001712), the LOCATE Biobank includes human coronary arteries (right coronary artery, left anterior descending artery, circumflex artery) excised postmortem during routine autopsy, preserved in 4% formaldehyde, embedded in paraffin, and cut into 5-µm sections. To assess the correlation between macrophage EHD1 expression and plaque stage, cross-sections of human coronary arteries were subjected to immunofluorescence staining with EHD1 and CD68 antibodies. Images were acquired using a Zeiss fluorescence microscope, and the mean fluorescence intensity (MFI) of EHD1 in CD68⁺ cells was quantified using ImageJ software.

### Statistical analysis

All results are presented as mean ± SEM. Statistical significance was determined using GraphPad Prism software. P values were calculated using the Student’s t-test for normally distributed data and the Mann-Whitney rank sum test for non-normally distributed data. One-way ANOVA with Dunnett post-test was used to analyze multiple groups with only one variable test. The Pearson’s correlation coefficient and p-value were determined by R (v4.3.2).

### Data and code availability

The RNA-seq data has been deposited at the Gene Expression Omnibus (GEO). The deposited data will be publicly available as of the date of publication. This paper did not report original code. Any additional information required to reanalyze the data reported in this paper is available from the lead contact upon request.

Detailed experimental methods and the Major Resources Table are available in the Supplemental Material.

## Results

### EHD1 expression in macrophages is induced in mouse and human atherosclerotic plaques

Macrophages play an essential role in atherosclerosis through their capability for phagocytosis (a form of endocytosis) and cargo recycling,^30^ however, the specific mechanisms underlying these processes are not completely understood. EHD-family proteins are well-known to regulate endocytosis. However, their role in atherosclerosis has not been characterized. Therefore, we analyzed the expression of EHD proteins in aortic macrophages by processing a published scRNA-seq dataset where the transcriptome of sorted CD45^+^ cells from whole aortas of *Ldlr^-/-^* mice fed WD was profiled.^31^ Consistent with the published results, we were able to identify 11 leukocyte clusters (**Figure S1A**). Among them, 8 clusters (clusters 0–5, 7, and 8) were macrophages based on their high-level expression of macrophage marker F4/80 (*Adgre1*) (**Figure S1B**). We then analyzed the expression profiles for all EHD proteins. Interestingly, we found that only *Ehd1* and *Ehd4* were expressed in aortic macrophages (**Figure S1C**). Unlike *Ehd4*, which was universally expressed in all macrophage clusters, *Ehd1* was enriched in macrophage clusters 2 and 5, with enriched genes involved in regulating inflammatory response, apoptosis, NF-kB activity, and Toll-like receptor 4 (TLR4) signaling (**Figure S1D**). To determine whether EHD1 expression in macrophages is altered during the progression of atherosclerosis, we compared EHD1 expression in lesional macrophages from *Ldlr^-/-^* mice fed WD for 7 weeks (early lesions) *vs.* 16 weeks (advanced lesions) and found that macrophages in advanced lesions expressed significantly more EHD1 than those in early lesions (**Figure 1A**). We also examined human coronary arteries and compared early lesions (Stage II) *vs.* advanced lesions (Stage IV). We found that EHD1 was induced in advanced lesions in humans as well (**Figure 1B**). We then analyzed gene expression in macrophages from the scRNA-seq of human atherosclerotic plaques,^32^ and found that the expression of *EHD1* was positively correlated with the expression of the pro-inflammatory genes *TNF* and *IL1B* (**Figure 1C and 1D**). This data suggests that EHD1 may be involved in certain cellular processes that cause chronic inflammation and promote plaque progression. We next sought to determine the possible mechanisms of induced EHD1 expression in macrophages. As EHD1^+^ aortic macrophages have enriched genes involving TLR4 signaling (**Figure S1D**), and TLR4 is likely activated in lesions by endogenous ligands and contributes to the progression of atherosclerosis,^33,34^ we activated TLR4 using LPS in BMDMs transfected with either scrambled siRNA (siSrc) or *Ehd1* siRNA (siEhd1). We found that LPS dramatically induced EHD1 expression at both the protein and mRNA levels (**Figure 1E and 1F**). Interestingly, LPS-induced EHD1 expression partially depended on the transcription factor hypoxia-inducible factor 1 alpha (HIF1α) (**Figure S1E and S1F**). We also treated BMDMs with atherogenic oxLDL and found that oxLDL induced EHD1 expression at protein but not mRNA levels (**Figure 1G and 1H**). This data indicates that atherogenic stimuli can induce EHD1 in macrophages, although this may occur through different molecular mechanisms.

**Figure 1.**
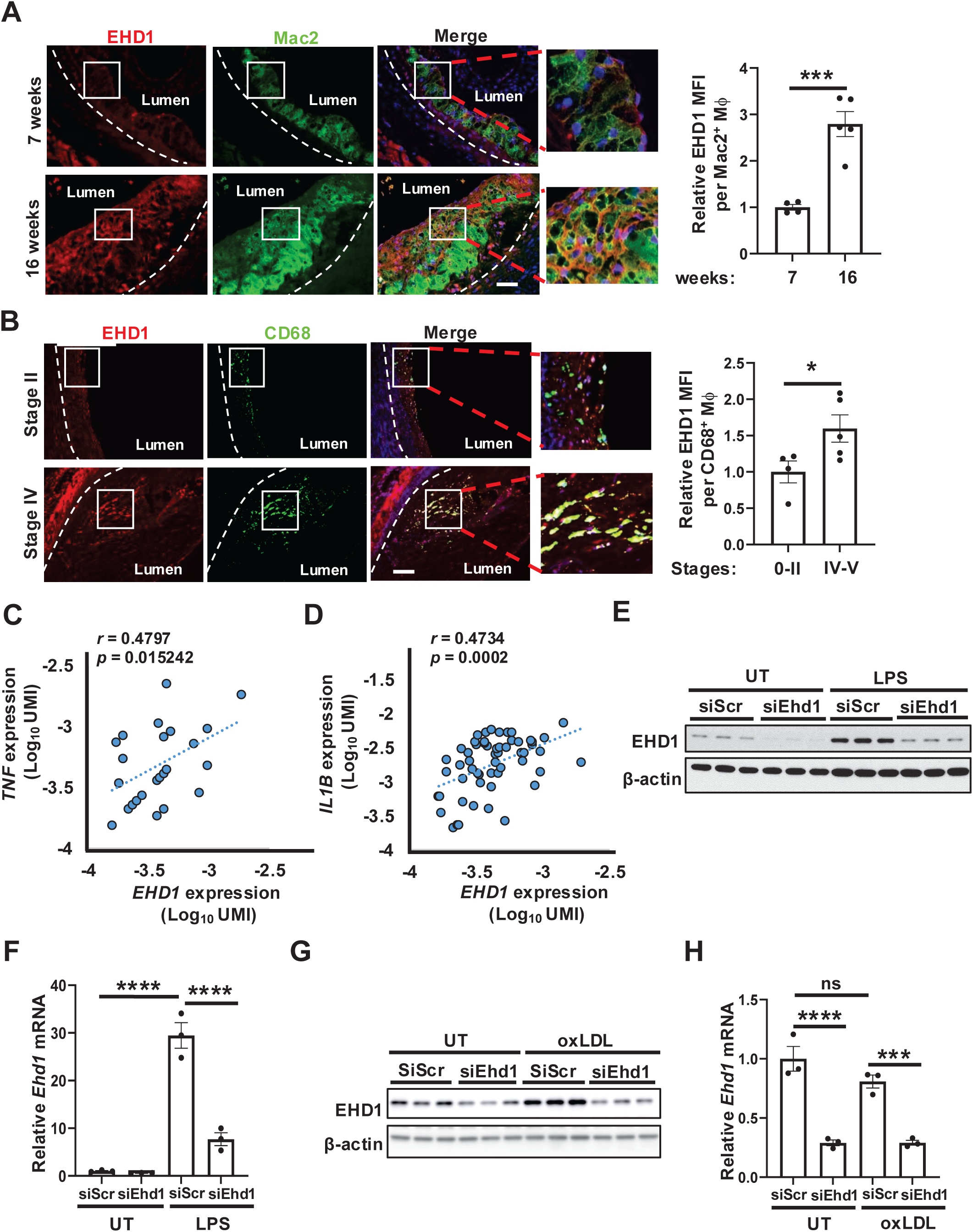
EHD1 expression in macrophages is induced in mouse and human atherosclerotic plaques. **A.** Aortic root sections from atherosclerotic male *Ldlr^−/−^*mice were co-stained for EHD1, Mac2, and DAPI. The mean fluorescence intensity (MFI) of EHD1 in Mac2^+^ cells was quantified with ImageJ. Representative images are presented (*n* = 4–5 /group). Scale bar, 100 µm. **B.** Human coronary artery sections were co-stained for EHD1, CD68, and DAPI. The MFI of EHD1 in CD68^+^ cells was quantified with ImageJ. Representative images are presented (*n* = 4–5/group). Scale bar, 100 µm. **C–D.** The scatter plots showed expression of *EHD1* (log_10_ UMI value) with *TNF* (C) and *IL1B* (D) in macrophages from human atherosclerotic plaques. The linear regression line, correlation coefficient (r, Pearson), and p-value in each plot is shown. **E–F.** BMDMs were treated with either scrambled control siRNA (siSrc) or siEhd1, followed by the treatment of 100 ng/mL LPS for 24 h. Protein (E) and mRNA (F) expression levels of EHD1 were determined by immunoblotting and qPCR, respectively. **G–H.** Control or EHD1-silenced cells were treated with 50 ug/mL oxLDL for 48 h, followed by EHD1 protein (G) and mRNA (H) measurement (*n* = 3 biological replicates). Data is presented as mean ± SEM. Data was analyzed using Student’s t-test in A and B, and one-way ANOVA followed by Dunnett’s multiple comparisons test in F and H. *p < 0.05, ***p < 0.001, ****p < 0.0001. n.s., not significant.

### Hematopoietic EHD1 deletion suppresses plaque progression and inflammation

To explore whether there might be a causal role of EHD1 in atherosclerosis, we performed a bone marrow transplantation experiment by transplanting either wild type (WT) or *Ehd1^-/-^* bone marrow cells into irradiated *Ldlr^-/-^* mice. Note that EHD1 deficiency did not compensatorily enhance EHD4 expression in macrophages (**Figure S2A**). After 6 weeks post-transplantation, these mice were fed with a WD for 12 weeks, a time point when the aortic root plaques have features of both early and advanced lesions. Although there were no differences in body weight, plasma cholesterol, or triglycerides (TGs) between the two groups of atherosclerotic mice (**Figure S2B–S2D**), the lesion size in H&E-stained aortic roots was significantly decreased in *Ehd1^-/-^*compared with WT bone marrow–transplanted mice (**Figure 2A**). Consistent with a decreased lesion size, Mac2^+^ macrophage staining area was decreased in *Ehd1^-/-^* plaques (**Figure 2B**). However, when we examined plaque necrosis by measuring H&E-negative acellular areas (≥ 3,000 mm^2^) in the intima, as we previously published,^29,35^ we observed that necrotic core areas were similar in these two groups (**Figure S2E**). Moreover, there was no significant difference in collagen cap thickness as determined by picrosirius red staining (**Figure S2F)**. These results suggest that EHD1 enhances lesion size but does not affect plaque stability.

**Figure 2.**
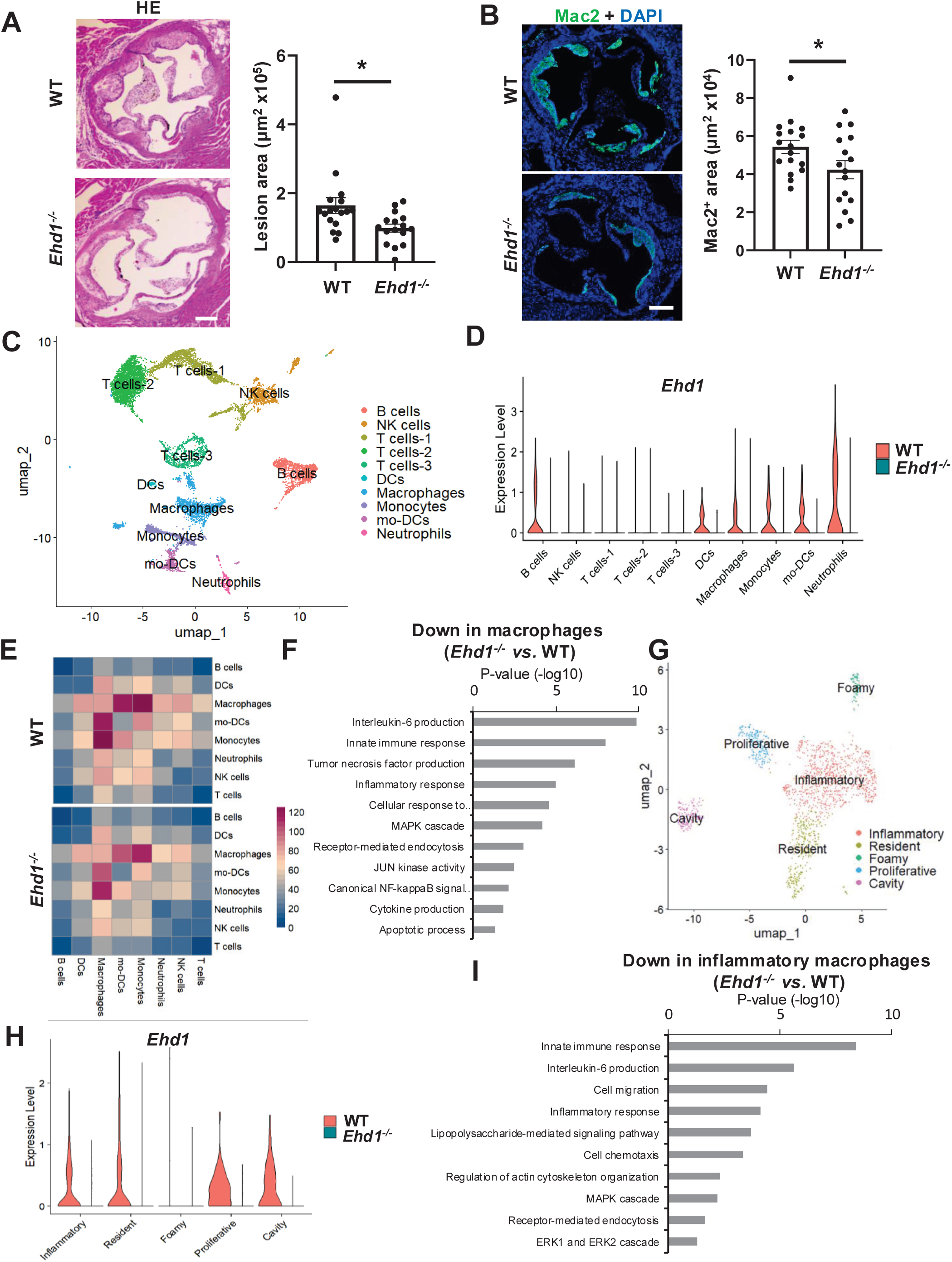
Hematopoietic EHD1 deficiency suppresses plaque progression and inflammation. **A.** Representative H&E-stained aortic root lesions and quantification of total lesion area from male *Ldlr^−/−^* recipients transplanted with WT or *Ehd1^-/-^* bone marrow cells after 12 weeks of WD feeding. Lesion area was quantified with ImageJ (*n* = 16–17/group). Scale bar, 200 μm. **B.** Representative immunofluorescence images of Mac2 and DAPI of aortic root lesions and quantification of total Mac2^+^ area by ImageJ (*n* = 16–17/group). Scale bar, 200 μm. **C.** UMAP visualization of CD45^+^ cells isolated from male *Ldlr^−/−^* recipient aorta lesions. Each annotated cell type is indicated with a different color. **D.** Violin plots showing *Ehd1* expression in different immune cell types from WT or *Ehd1^-/-^* aorta. **E.** CellphoneDB cell–cell interaction heatmap between different cell types in WT or *Ehd1^-/-^* aorta CD45^+^ cells. A color scale indicates the interaction counts. **F.** GO analysis for the downregulated DEGs in macrophages. **G.** UMAP visualization of different aortic macrophage subtypes. **H.** Violin plots showing *Ehd1* expression in each macrophage subtypes from WT or *Ehd1^-/-^*aorta. **I.** GO analysis for the downregulated DEGs in inflammatory macrophages isolated from *Ehd1^-/-^* vs. WT aorta. Data is presented as mean ± SEM. *p < 0.05 by Student’s t-test.

To determine how EHD1 alters the immune microenvironment in atherosclerotic plaques, we performed scRNA-seq on pooled CD45^+^ aortic cells from irradiated *Ldlr^-/-^* mice transplanted with either WT or *Ehd1^-/-^* bone marrow cells. A total of 10,405 single-cell transcriptomes were analyzed. Clustering visualized on uniform manifold approximation and projection (UMAP) revealed ten major clusters of cells, including B cells, NK cells, T cells, dendritic cells, macrophages, monocytes, and neutrophils in both WT and *Ehd1^-/-^*aortas (**Figures 2C, S3A and S3B**). Interestingly, we observed that B cells, dendritic cells, and neutrophiles also expressed EHD1 in atherosclerotic aorta (**Figure 2D).** Although EHD1 deletion did not alter macrophage composition among immune cells (**Figure S3C**), CellphoneDB analysis demonstrated that the overall interactions of macrophages with other immune cells, including neutrophils, NK cells, and T cells, were attenuated (**Figure 2E**). We next compared differentially expressed genes (DEGs) from psudo bulk analysis between WT and *Ehd1^-/-^* aortic macrophages. We identified 310 downregulated and 39 upregulated genes in *Ehd1^-/-^* compared with WT aortic macrophages (**Figure S3D**). Gene ontology (GO) analysis for these DEGs demonstrated that biological processes including IL6 production, TNFα production, NF-kB signaling, and apoptosis were suppressed in *Ehd1^-/-^* aortic macrophages (**Figure 2F**), while protein localization to endoplasmic reticulum (ER), catabolic process, and calcium signaling were upregulated (**Figure S3E**). We further subclustered macrophages into five major populations—inflammatory, resistant, foamy, proliferative, and cavity macrophages (**Figures 2G, S4A and S4B**). EHD1 was expressed in all the macrophage subclusters except foamy cells (**Figure 2H**) and its deletion did not alter the composition of macrophage subpopulations (**Figure S4C**). Because the inflammatory macrophage was the predominant macrophage subcluster, we then analyzed DEGs between WT and *Ehd1^-/-^* inflammatory macrophages. We identified 129 downregulated and 31 upregulated genes in *Ehd1^-/-^* as compared with WT inflammatory macrophages (**Figure S4D**). In line with the decreased inflammatory response observed in *Ehd1^-/-^* total aortic macrophages, the GO analysis for these DEGs demonstrated that innate immune response, IL-6 production, and LPS-mediated signaling were suppressed in *Ehd1^-/-^* inflammatory macrophages (**Figure 2I**), while pathways involving protein catabolic process, calcium signaling, and protein palmitoylation were upregulated (**Figure S4E**). This data indicates that EHD1 promotes inflammation in atherosclerotic aorta.

### EHD1 deficiency attenuates NF-kB-mediated inflammation in macrophages

To determine whether EHD1 directly regulates inflammatory response in macrophages, we silenced EHD1 in BMDMs and performed RNA-seq to explore its effect on inflammation. In unstimulated cells, we identified 630 downregulated and 879 upregulated DEGs by comparing siEhd1-with siSrc-treated cells (**Figure S5A**). The GO analysis for the DEGs indicated that pathways including NF-kB signaling, TNF-α production, IL-6 production, and apoptosis were attenuated while cell cycle and protein modification were upregulated in EHD1-silenced macrophages (**Figure S5A**). Similarly, in LPS-treated macrophages, the inflammatory response was suppressed while cell cycle was induced upon EHD1 deficiency (**Figures 3A and S5B**). We then verified the effect of EHD1 on the NF-kB-mediated inflammation pathway by measuring NF-kB activity and cytokine production. As expected, LPS induced the phosphorylation of p65, an essential subunit of NF-kB transcription factor complex, indicating an increase in NF-kB activity. However, the LPS-induced NF-kB activation was diminished in both EHD1-silenced and knockout macrophages (**Figure 3B and 3C**). In line with the decreased NF-kB activity, the pro-inflammatory mediators, including NOD-, LRR-, and pyrin domain-containing protein 3 (NLRP3), IL-1β, TNF-α, and IL-6 were reduced at both protein and mRNA levels in EHD1-deficient macrophages (**Figure 3B-E; Figure S5C and S5D**). To determine whether these *in vitro* studies are recapitulated *in vivo*, we analyzed the pro-inflammatory mediators in aortic inflammatory macrophages and found that the expression levels of *Il1b*, *Tnf*, and *Nlrp3* were decreased in *Ehd1^-/-^* aorta (**Figure 3F**). The decreased IL-1β expression was also confirmed in *Ehd1^-/-^* aortic macrophages (**Figure 3G**). Consistent with the concept that excessive inflammatory cytokines, including IL-1β, can induce ROS and activate apoptotic pathways,^36–40^ and supported by RNA-seq data showing that EHD1 deficiency suppresses apoptosis, EHD1 deficiency eliminated LPS-induced ROS and apoptosis, as indicated by decreased DAPI^+^ cells and decreased cleaved caspase 3 (**Figure 3H–J**). The decreased IL-1β and apoptosis were also observed in oxLDL-treated EHD1-silenced cells (**Figure S5E**). In line with these *in vitro* findings, *Ehd1^-/-^*plaques had decreased cell death (**Figure 3K**). This data suggests that EHD1 promotes atherosclerosis potentially through activating NF-kB-mediated inflammation.

**Figure 3.**
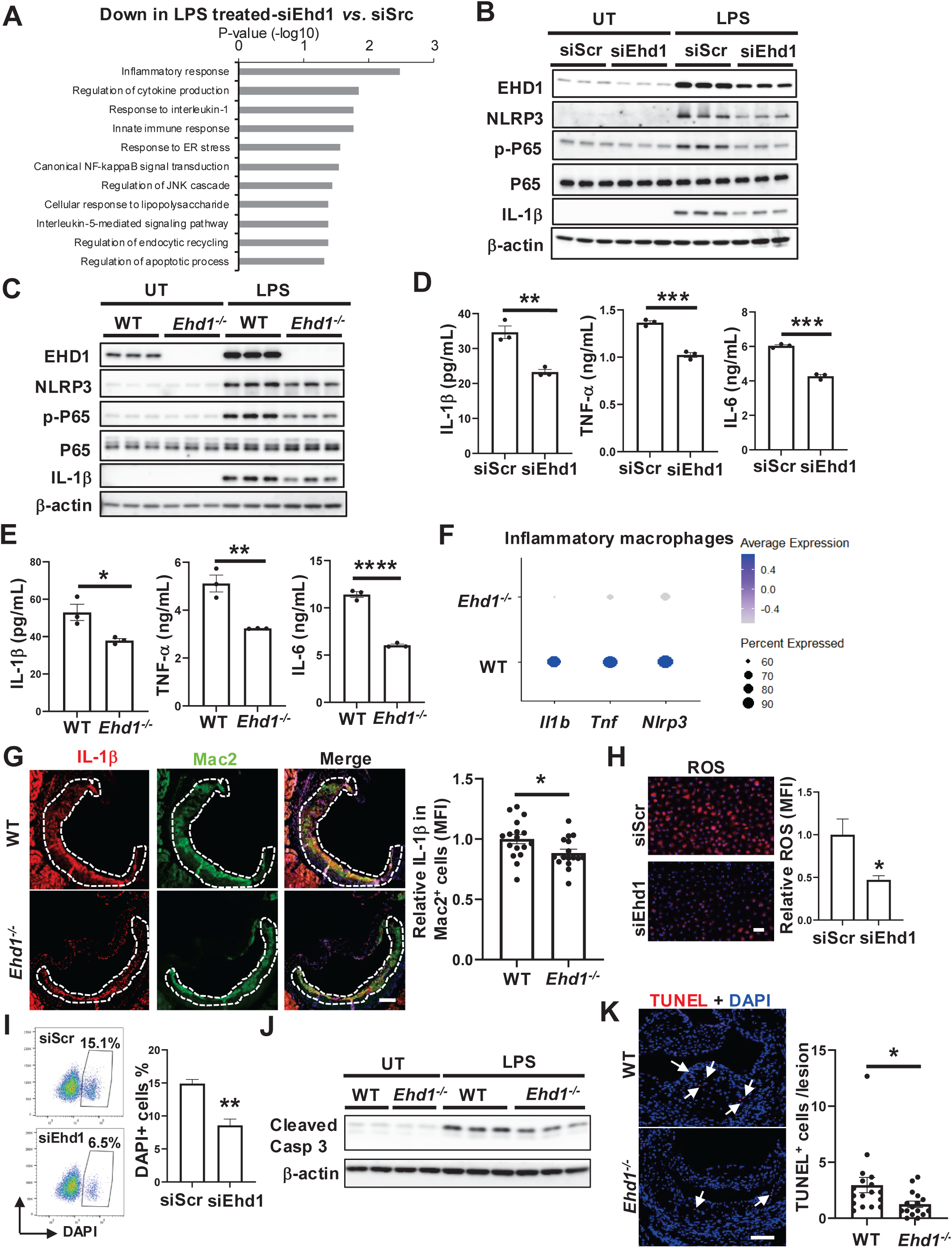
EHD1 deficiency attenuates NF-kB-mediated inflammation in macrophages. **A.** GO analysis for the downregulated DEGs in BMDMs transfected with siRNAs, followed by treatment with 100 ng/mL LPS for 18 h. **B–C.** Immunoblots of EHD1, NLRP3, p-P65, P65, and IL1β from siEhd1-treated (B) and *Ehd1^-/-^* (C) cells upon LPS treatment. β-actin was used as the loading control. **D–E.** Secreted IL1β, TNFa, and IL6 levels in culture media from siEhd1-treated (D) and *Ehd1^-/-^* (E) cells were assayed by ELISA (*n* = 3 biological replicates). **F.** Dot plots showing *Il1b*, *Tnf*, and *Nlrp3* expressions in inflammatory macrophages. A color scale of average expression is shown. **G.** Representative immunofluorescence images of IL1β, Mac2, and DAPI in aortic root lesions, and quantification of IL1β MFI in Mac2^+^ cells by ImageJ (*n* = 16–17/group). Scale bar, 100 μm. **H.** Representative ROS images from LPS-treated EHD1-silenced cells and quantification of ROS MFI (*n* = 3 biological replicates). Scale bar, 100 µm. **I.** Representative flow cytometry plots of DAPI⁺ BMDMs with corresponding quantification (*n* = 3 biological replicates). **J.** Immunoblots of Cleaved Caspase 3 (Casp 3) in BMDMs treated with LPS for 24 h. **K.** Representative immunofluorescence images of TUNEL and DAPI of aortic root lesions, and quantification of TUNEL^+^ cells per lesion (*n* = 16–17/group). Arrows indicate TUNEL^+^ cells. Scale bar, 100 μm. Data is presented as mean ± SEM. *p < 0.05, **p < 0.01, ***p < 0.001, ****p < 0.0001 by Student’s t-test.

### EHD1 deficiency suppresses endocytic recycling of TNFR2

We next sought to determine the mechanism by which EHD1, as an endocytic recycling regulator, promotes NF-kB activity. We assayed the cell-surface levels of Cluster of Differentiation 36 (CD36), TLR4, and TNFR2 that have been shown to activate NF-kB signaling^41–43^ on macrophages by flow cytometry. Strikingly, we found that EHD1 deficiency specifically reduced cell-surface TNFR2 but not CD36 and TLR4 (**Figure 4A and 4B; Figure S5F and S5G**). Note that EHD1 deficiency did not alter *Tnfr2* mRNA expression, indicating that the reduced cell-surface TNFR2 is not due to decreased expression (**Figure S5H and S5I**). We then performed a pulse-chase experiment to assay the effect of EHD1 silencing on endocytic recycling of TNFR2. We conducted an antibody-triggered endocytosis assay^44,45^ by incubating macrophages with anti-TNFR2 antibodies. We incubated cells with anti-TNFR2 antibodies on ice to allow antibodies to bind to TNFR2 on the cell surface. Cells were then placed at 37°C to trigger TNFR2 internalization (pulse). After acid-stripping to remove uninternalized TNFR2-TNFR2 antibody complex, cells were incubated in complete media to allow internalized TNFR2 to recycle back to the cell surface (chase), followed by a second acid-stripping to remove the recycled TNFR2 on the cell surface. The remaining intracellular TNFR2 was then visualized by confocal microscopy. As expected, we observed that the remaining TNFR2 was decreased during recycling chase phases in control cells, but the intracellular TNFR2 was sustained in EHD1-silenced cells, indicating that EHD1 deficiency reduces TNFR2 recycling (**Figure 4C**). We found that TNFR2 intensity was reduced in *Ehd1^-/-^* aortic macrophages (**Figure 4D**), which is consistent with the concept that impaired recycling leads to degradation.^46,47^ This data indicates that EHD1 promotes NF-kB-mediated inflammation potentially through enhancing TNFR2 recycling to the cell surface.

**Figure 4.**
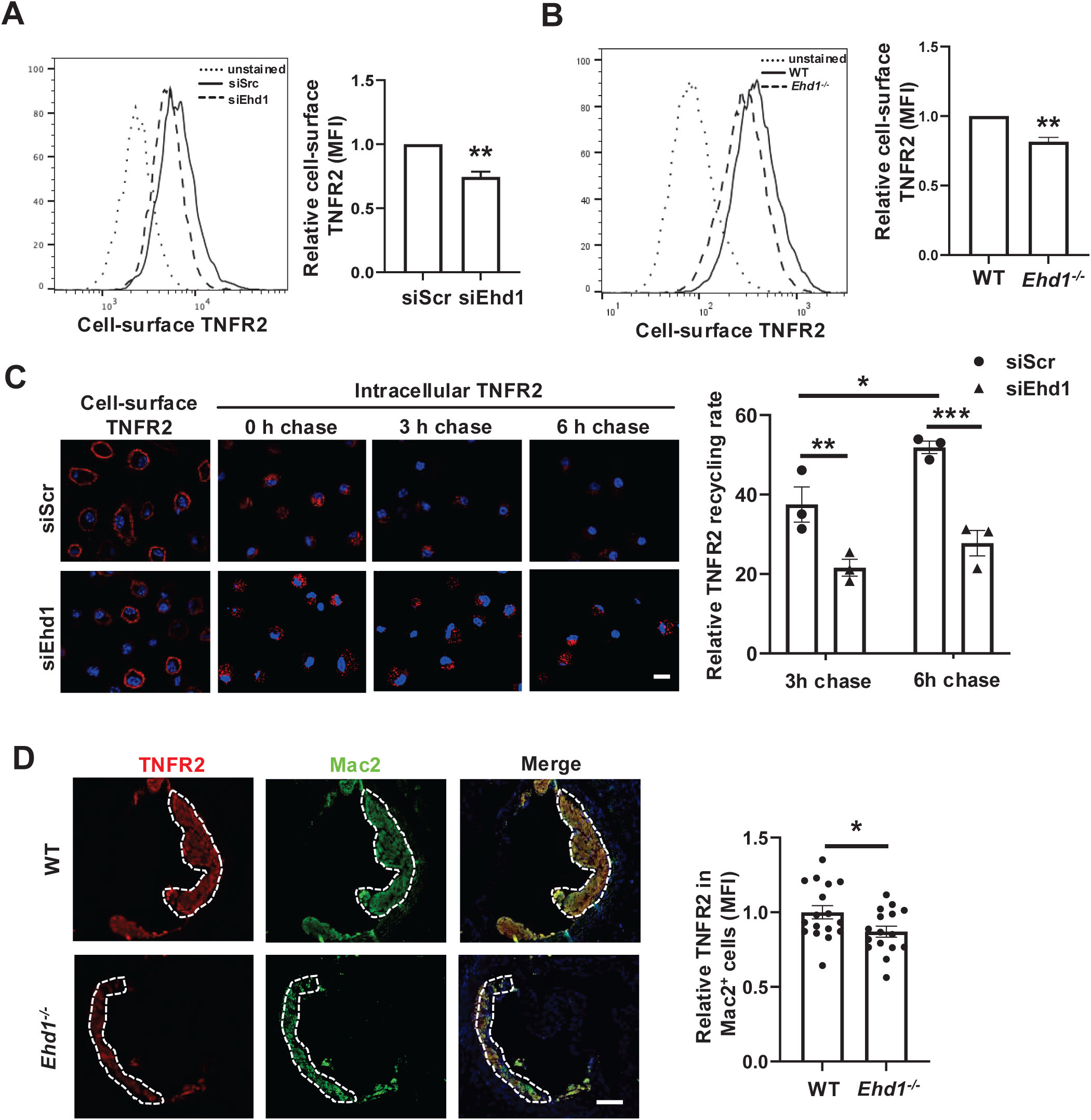
EHD1 deficiency suppresses endocytic recycling of TNFR2. A–B. Cell-surface levels of TNFR2 in siEhd1-treated (A) and *Ehd1^-/-^* (B) BMDMs were detected by flow cytometry (*n* = 3 biological replicates). **C.** siRNA-treated BMDMs were incubated with anti-TNFR2 antibodies on ice to detect cell-surface TNFR2. Cells were placed at 37°C to allow TNFR2 internalization, followed by acid-stripping and chasing in complete media to allow TNFR2 recycling. After a second acid-stripping, the remaining intracellular TNFR2 was visualized by confocal microscopy. The recycling rate of TNFR2 was calculated as (total internalized TNFR2 − remaining TNFR2) / total internalized TNFR2 (*n* = 3 biological replicates). Scale bar, 10 µm. **D.** Representative immunofluorescence images of TNFR2, Mac2, and DAPI in aortic root lesions, and quantification of TNFR2 MFI in Mac2^+^ cells by ImageJ (*n* = 16–17/group). Scale bar, 100 μm. Data is presented as mean ± SEM. Data was analyzed using Student’s t-test in A, B, and D, and two-way ANOVA followed by uncorrected Fisher’s LSD test in C. *p < 0.05, **p < 0.01, ***p < 0.001.

### EHD1 interacts with and stabilizes sortilin

In addition to endocytic recycling, EHD1 has been shown to promote retromer-mediated retrograde transport from endosomes to trans Golgi (TGN), a cellular process of rescuing cargo from lysosomal degradation.^18,19^ Retromer is a large protein complex composed of vacuolar protein sorting (VPS) 26, VPS29, and VPS35, and plays a crucial role in stabilizing sortilin, a risk factor for cardiovascular disease in humans.^48^ As sortilin has been shown to promote atherosclerosis by inducing IL-6 secretion in macrophages,^28^ and our scRNA-seq data showed that EHD1 deficiency suppressed IL-6, we proposed that EHD1 stabilizes sortilin by promoting retromer-mediated retrograde transport, thus leading to increased IL-6 and atherosclerosis progression. We first observed that EHD1, sortilin, and VPS26α are in the same protein complex in macrophages by performing a co-immunoprecipitation assay (**Figure 5A**). We then silenced EHD1 and found that EHD1 deficiency significantly reduced sortilin protein but not mRNA levels in untreated and oxLDL-treated macrophages (**Figures 5B and S5J**). Similar results showing reduced sortilin were also obtained in *Ehd1^-/-^* macrophages (**Figure 5C**). Consistent with these *in vitro* studies, sortilin levels in Mac2^+^ macrophages were decreased in *Ehd1^-/-^* lesions (**Figure 5D**). This data suggests that EHD1-promoted sortilin stability may contribute to the progression of atherosclerosis.

**Figure 5.**
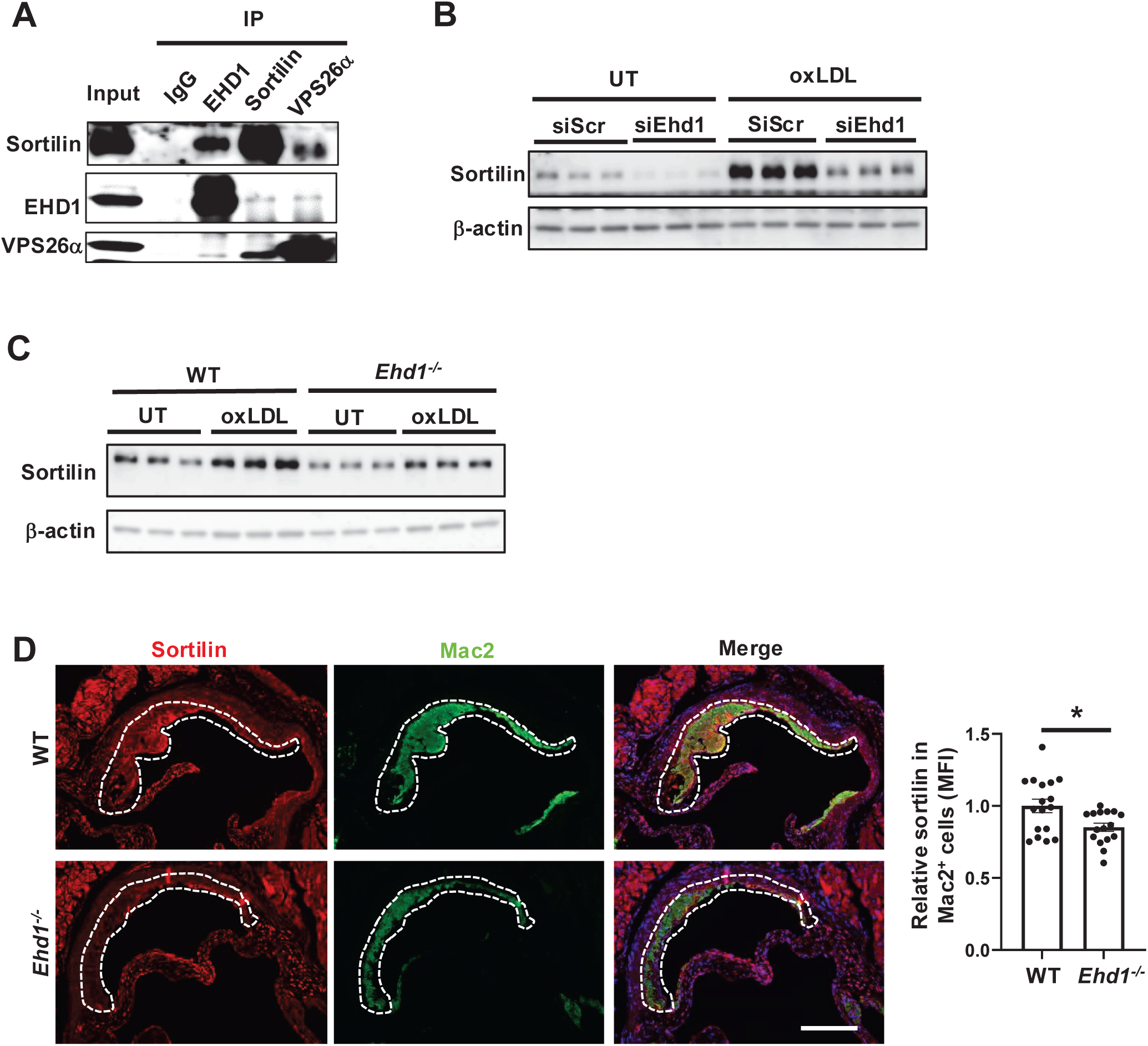
EHD1 interacts with and stabilizes sortilin. **A.** Co-immunoprecipitation of EHD1, sortilin, and VPS26α in BMDMs. **B.** Immunoblots of sortilin in BMDMs transfected with siRNAs, followed by treatment with 50 µg/mL oxLDL for 48 h. **C.** Immunoblots of sortilin in WT and *Ehd1^- /-^* BMDMs, followed by oxLDL treatment. **D.** Representative immunofluorescence images of sortilin, Mac2, and DAPI in aortic root lesions, and quantification of sortilin MFI in Mac2^+^ cells by ImageJ (*n* = 16–17/group). Scale bar, 100 μm. Data is presented as mean ± SEM. *p < 0.05 by Student’s t-test.

## Discussion

In this study, we demonstrated new roles of membrane trafficking involving endocytic recycling and retrograde transport in regulating the pro-inflammatory response in macrophages. Specifically, we found that EHD1-mediated endocytic recycling of TNFR2 and EHD1-stabilized sortilin, a retrograde cargo, promotes inflammation in macrophages, leading to accelerated atherosclerosis (**Figure S6A**). Among all the EHD proteins, only EHD1 and EHD4 are detectable in aortic macrophages. However, the expression patterns of EHD1 and EHD4 are quite different. EHD1 is enriched in pro-inflammatory macrophages, while EHD4 seems to be expressed ubiquitously in all aortic macrophages. Therefore, whether macrophage EHD4 plays a role in atherosclerosis through a mechanism different from EHD1 requires future investigation.

Our sc-RNA seq showed that EHD1 is not expressed in foamy cells in aorta, consistent with lipid-poor macrophages that exhibit a higher inflammatory profile.^7^ Our study for the first time revealed that EHD1-mediated membrane trafficking regulates the activation of NF-kB, the master transcription factor governing inflammatory responses. There are many cell-surface receptors that can signal and activate NF-kB in macrophages.^43,49^ Interestingly, we found that EHD1 specifically maintains the cell-surface TNFR2 rather than CD36 and TLR4. Cell-surface TNFR2 activates NF-kB via TRAF2 and induces the expression of cytokines, including TNF-α, which engages with TNFR2, leading to an amplification of inflammatory responses.^43,50^ Therefore, our study indicates that EHD1 is essential for sustaining the positive feedback loop of inflammatory responses by promoting the endocytic recycling of TNFR2 to the cell surface in macrophages. Future studies are needed to determine the sorting motifs in the TNFR2 needed for EHD1-mediated endocytic recycling. In addition to the well-established role in receptor recycling, EHD1 has been shown to interact with the retromer complex to regulate retrograde transport of cargoes from endosomes to TGN in HeLa cells.^18^ Of note, sortilin can undergo retrograde transport through the retromer complex and escape from lysosomal degradation.^51,52^ Along this line, we found that EHD1 interacts with retromer and sortilin, and that EHD1 stabilizes sortilin in macrophages. Human genome-wide association studies (GWAS) have revealed a strong association between sortilin and cardiovascular diseases, including atherosclerosis.^53,54^ Emerging evidence shows that sortilin drives the pathogenesis of atherosclerosis by serving as a key receptor for cytokines and lipids.^28,55–57^ Based on these human genetic studies of sortilin, our study highlights the human relevance of EHD1 in atherosclerosis through its link to sortilin. However, the cellular mechanisms by which EHD1 regulates retromer-mediated retrograde transport of sortilin require future exploration. For instance, it will be interesting to determine whether EHD1 controls the stability and the intracellular distribution of retromer complex, and to elucidate whether the ATPase activity of EHD1 is required for the retrograde transport of sortilin.

Previous studies have shown that EHD proteins modulate gap junctions and ion channels in cardiomyocytes.^26,27^ However, the roles of EHD proteins in immune responses have never been explored. Our study for the first time established the essential role of EHD1 in regulating macrophage-mediated inflammatory responses in the context of atherosclerosis. In addition to macrophages, we saw that EHD1 is expressed in B cells, dendritic cells (DCs), and neutrophils in the aorta. Given that endocytic membrane trafficking has been shown to regulate antigen procession and cell polarization through modulating cell-surface receptors in B cells, DCs, and neutrophils,^58–60^ future studies are needed to elucidate the roles of EHD protein–mediated membrane trafficking in those immune cells. Of note, our scRNA-seq showed that EHD1 deletion attenuates the communication of macrophages with other immune cells. It will be worthwhile to dissect the ligand and receptor pairs in different immune cell populations that are mediated by EHD1 and determine whether those interactions contribute to the pathogenesis of atherosclerosis.

Endocytic membrane trafficking is pivotal for cell physiology, as receptor internalization and recycling are fundamental to ensure cellular sensing and division, cell migration, and cell–cell communication.^61^ Dysregulation of endocytic membrane trafficking has consistently been found to correlate with human diseases, including cancers and neurodegeneration.^10,62,63^ In our study, we found that EHD1 expression is induced in both human and mouse atherosclerotic plaques, indicating a dysregulation of endocytic membrane trafficking in the pathogenesis of atherosclerosis. Interestingly, we found that LPS-induced EHD1 expression is partially blunted by HIF1α silencing. Given that HIF1α has been shown to promote atherosclerosis,^64^ whether EHD1 induction requires the activation of HIF1α during the progression of atherosclerosis needs to be determined in the future.

As professional phagocytes, macrophages play major roles in clearing apoptotic cells (ACs), a cellular process called “efferocytosis,” in atherosclerotic plaques.^65^ When efferocytosis is impaired, uncleared ACs undergo post-apoptotic necrosis and release immunogenic epitopes and pro-inflammatory mediators, leading to chronic inflammation.^29,65^ Similar to endocytosis, efferocytosis involves cytoskeleton rearrangement and the intracellular transport of cargo-containing vesicles. However, the role of endocytic regulators in the cellular transport of AC-derived cargo during efferocytosis is unknown. Therefore, there is a need to study whether EHD proteins regulate the cell-surface levels of efferocytosis receptors such as MerTK, LRP1, and Tim4 that are essential for AC internalization,^65^ and to determine whether endosomal membrane trafficking controls AC digestion upon engulfment.

In summary, we for the first time explored EHD protein–mediated membrane trafficking in macrophage biology in the context of atherosclerosis. We analyzed the expression patterns of EHD proteins in different aortic immune cells by scRNA-seq of mouse and human plaques. Our study also highlights the novel regulation of EHD1 in stabilizing sortilin, which has been identified as a risk factor for atherosclerosis in human GWAS.

## Acknowledgments

The authors thank the Single Cell Analysis Core and Columbia Genome Center at Columbia University for providing services related to the generation of RNA-sequencing data. The scheme was generated with BioRender.

## Author contributions

B.C. and F.M. developed the study concept and experimental design; F.M., Y.L., N.G., and B.C. conducted all the *in vivo* and *in vitro* experiments; Y.X., J.C.K., and W.L. helped with atherosclerosis studies; T.F. and Y.Z. assisted with aortic cell isolation for scRNA-seq; L.X. and X.H. helped analyze RNA sequencing data; J.G.T., A.Y.J., A.W.O., and O.R. provided human coronary artery specimens and critical review; B.C. and F.M. interpreted the data and wrote the manuscript; and all the coauthors participated in the manuscript editing.

## Nonstandard abbreviations and acronyms

EHD: c-terminal Eps15 Homology Domain
BMDM: bone marrow–derived macrophages
LPS: lipopolysaccharide
oxLDL: oxidized low-density lipoprotein
NF-kB: nuclear factor kappa B
NLRP3: NOD-, LRR-, and pyrin domain–containing protein 3
TNF-α: tumor necrosis factor-alpha
IL-1β: interleukin-1 beta
IL-6: interleukin-6
TUNEL: terminal deoxynucleotidyl transferase dUTP nick-end labeling
ROS: reactive oxygen species
HIF: hypoxia-inducible factor 1 alpha
TNFR2: tumor necrosis factor receptor 2
CD36: cluster of differentiation 36
TLR4: Toll-like receptor 4
VPS26α: vacuolar protein sorting-associated protein 26A
WD: Western-type diet
DEG: differentially expressed genes
GO: gene ontology

## SUPPLEMENTAL MATERIAL

### Detailed Methods

#### Atherosclerotic lesional analyses

After 12 weeks of WD, *Ldlr^−/−^* mice were fasted overnight and sacrificed with isoflurane. Body weights were recorded, and blood was collected by cardiac puncture. Following perfusion with 5 mL PBS, hearts were harvested and fixed in 4% PFA at 4°C overnight. The next day, tissues were transferred to 75% ethanol and submitted to the Mount Sinai Histology Core Facility for paraffin embedding. When the first lesion appeared, two sections of the aortic root (5-μm thickness) were collected onto a single slide using a microtome. For each mouse, every fifth slide was selected for hematoxylin and eosin (H&E) staining. Lesion size and necrotic core area (≥ 3,000 mm^2^) were quantified using ImageJ software. For fibrous cap thickness analysis, picrosirius red staining was performed on sections from comparable regions of the aortic root. In each section, the cap of the largest lesion was measured at 2-μm intervals, and the average thickness was reported.

#### Immunofluorescence staining and TUNEL assay

Paraffin sections were deparaffinized, rehydrated through graded ethanol (100–70%), and subjected to antigen retrieval in citric acid buffer using a pressure cooker for 10 min. After cooling and PBS washes, sections were blocked with 5% donkey serum for 1 h at room temperature (RT) and incubated overnight at 4°C with primary antibodies diluted in 1% donkey serum. The next day, sections were washed with PBS and incubated with secondary antibodies for 1 h at RT. Slides were mounted with anti-fade solution and imaged on a Zeiss fluorescence microscope. Data was processed using ImageJ software. For TUNEL staining, aortic root sections were stained for dead cells using the In Situ Cell Death Detection Kit, TMR Red. Data was presented as average of the number of TUNEL+ cells per lesion for each mouse.

#### BMDM preparation

BMDMs were generated from 8-week-old male C57BL/6J mice, as well as from *Ehd1^−/−^* mice and their sex-matched WT littermates. The mice were euthanized and muscles were carefully removed from the hindlimbs. Bone marrow was flushed from the hindlimbs bones into a 40 µm cell strainer placed over a 50 mL tube using a syringe with a 25-gauge needle. The marrow was gently triturated through the strainer and washed with 10 mL medium. Cells were centrifuged at 600 x g for 6 min at 4°C and differentiated for 5–7 days in DMEM medium supplemented with 10% FBS, 1% penicillin-streptomycin, and 20% L-929 fibroblast-conditioned medium.

#### siRNA-mediated gene silencing

After 7 days of differentiation, BMDMs were detached and seeded either into 24-well tissue culture plates for RNA and protein extraction or into non-issue culture plates for flow cytometry. Cells in each well were incubated for approximately 48 h with 0.5 mL of culture medium containing 1.5 µL Lipofectamine RNAiMAX and 20 pmol siRNAs. SMART pool siRNAs targeting mouse Ehd1 was obtained from Dharmacon. The target sequence of mouse Hif1a was GAUAUGUUUACUAAAGGACAAGUCA.

#### ELISA analysis of cytokines from LPS- and oxLDL-treated cells

BMDMs from C57BL/6J mice transfected with siRNAs or from *Ehd1^−/−^* mice and their sex-matched WT littermates were washed twice with PBS and incubated in serum-free medium containing either 100 ng/mL LPS for 24 h or 50 µg/mL oxLDL for 48 h. Culture supernatants from LPS-treated cells were collected, and levels of TNF-α, IL-1β, and IL-6 were assessed by ELISA according to the manufacturer’s instructions.

#### Flow cytometry analysis

After treatment, BMDMs were detached and incubated with Fc block in FACS buffer at 4°C for 15 min. Cells were then stained with antibodies (1:100 dilution) for 30 min at 4°C. After being washed twice with FACS buffer, samples were analyzed on a BD FACS Canto II flow cytometer, and the data was processed using FlowJo™ software.

#### ROS detection

BMDMs were grown on cover glasses and incubated with 5 µM CellROX reagent for 30 min at 37°C. Cells were washed three times with PBS, fixed with 4% PFA for 10 min at RT, and permeabilized with 0.1% Triton X-100 for 10 min at RT. Cover glasses were mounted onto slides with anti-fade solution and imaged using a Zeiss fluorescence microscope. The data was processed using ImageJ software.

#### Immunoblotting

BMDMs with different treatments were lysed with 2x Laemmli sample buffer following 5 min boiling in 100°C heat block. Proteins were separated by electrophoresis in 4–20% Tris gels and transferred to 0.45 μm nitrocellulose membranes. The membranes were blocked for 1 h at RT in Tris-buffered saline/0.1% Tween 20 (TBST) containing 5% (wt/vol) nonfat milk and then incubated with primary antibodies in intercept blocking buffer at 4°C overnight. The membranes were then incubated with the appropriate secondary antibodies coupled to horseradish peroxidase, and proteins were detected by the SuperSignal™ West pico plus chemiluminescent substrate.

#### Coimmunoprecipitation (Co-IP)

After 7 days of differentiation, BMDMs from two 100 mm culture dishes were lysed on ice for 30 min with 2 mL ice-cold co-IP buffer (25 mM Tris-HCl, pH 7.5; 150 mM NaCl; 1.5 mM MgCl₂; 1 mM EDTA; 0.5% NP-40; and protease inhibitor cocktail). Lysates were centrifuged at 10,000 x g for 10 min at 4°C, and the supernatant was precleared with 30 µL protein G agarose beads for 2 h at 4°C. The precleared supernatant was evenly divided into four tubes and incubated overnight at 4°C with gentle rotation with rabbit anti-EHD1 antibody, rabbit anti-VPS26α antibody, rabbit anti-sortilin antibody, or rabbit IgG (negative control). Samples were then incubated with 50 µL prewashed protein G beads for 2 h at 4°C. Beads were collected by centrifugation and washed five times with 500 µL ice-cold co-IP buffer. Proteins were eluted with 30 µL 2x Laemmli sample buffer and analyzed by immunoblotting.

#### Quantitative RT-qPCR

Total RNA was extracted from treated BMDMs using the RNeasy kit. The quality and concentration of the RNA were assessed by absorbance at 260 and 280 nm using an EPOCH spectrophotometer. cDNA was synthesized from 0.5–1 μg RNA using the High-Capacity cDNA Reverse Transcription kit. Quantitative RT-PCR was performed with a light cycler 480 II Real-Time PCR system using SYBR Green Master Mix. The primer sequences are listed in Table S1.

#### TNFR2 pulse-chase assay

BMDMs were seeded on cover glasses and treated with siRNAs for 48 h. Cells were incubated with anti-TNFR2 antibody for 1 h at 37°C to allow TNFR2 internalization (pulse), followed by 1-min acid strip (0.5 m NaCl, 0.5% acetic acid, pH 3.0) to remove noninternalized TNFR2. Cells were then either fixed to detect total internalized TNFR2 (pulse) or placed in complete media at 37°C for 3 h or 6 h to allow TNFR2 recycling (chase). Upon chase, cells were subjected to a second acid stripping to remove recycled TNFR2 on the cell surface and fixed. Intracellular TNFR2 was detected with Alexa Fluor 568–conjugated goat anti-rabbit antibody diluted in staining solution containing 0.2% saponin (w/v) and 0.5% BSA (w/v) in PBS. Cover glasses were mounted with anti-fade solution and imaged using a confocal microscope. The intensity of remaining TNFR2 after chase was quantified by ImageJ. Percentage of recycling was calculated as a ratio of (total internalized TNFR2 − remaining TNFR2) / total internalized TNFR2.

#### Plasma cholesterol and triglycerides measurement

Plasma cholesterol and triglycerides levels were assayed using a total cholesterol assay kit and a L-Type triglyceride assay kit, as listed in the major resources table section.

#### Bulk RNA sequencing and data analysis

Total RNA from BMDMs was extracted using the RNeasy Mini Kit. RNA-seq library construction was conducted at Innomics Inc. (San Jose, CA) with a standard polyA-enrichment protocol. The sequencing was performed on a DNBSEQ G400 sequencer (MGI-tech, San Jose, CA) and 150-bp paired-end reads were obtained. The bulk RNA-seq data was processed as described in our previously published paper.^66, 67^

#### scRNA-seq and data analysis

Aortic CD45^+^ cells were isolated as previously described.^68^ Briefly, aortas from male *Ldlr^-/-^*mice (n = 7) fed a high-fat diet for 12 weeks were pooled and digested with an enzyme mixture. Aortic single cells were stained with anti-CD45 antibody and DAPI. DAPI-CD45^+^ cells were sorted by FACS and collected for scRNA-seq on the 10X Genomics Chromium platform at the Columbia Genome Center, Columbia University. scRNA-seq reads were aligned to the reference genome mm10-2020-A with the Cell Ranger program (v6.1.2, 10X Genomics). scRNA-seq data was processed as in our previously published paper.^1^ Briefly, the scRNA-seq filtered data matrices were imported in Seurat (v5.0.1) using R (v4.3.2). Cells with percentages of mitochondrial mRNAs over 10% were removed. The filtered data was processed by Seurat with the standard pipeline, including the NormalizeData, FindVariableFeatures, FindIntegrationAnchors, and IntegrateData functions, followed by the ScaleData, RunPCA, FindNeighbours, FindClusters, RunUMAP, and RunTSNE functions. If not specified, default settings were followed. We tested resolution values from 0.3 to 0.9 in the FindClusters function and determined 0.5 as the best resolution value. Cells were annotated based on marker genes identified from the FindMarkers function and the known lineage markers from the literature. The UMAP plot, heatmap, feature plot, dot plot, and violin plot with cell annotations were virtualized using Seurat. DEGs between the WT and *Ehd1^-/-^* groups were determined by p<0.05 and percent of expression >0.25 in both groups. GO analysis was performed on these DEGs using the DAVID tool (https://david.ncifcrf.gov/tools.jsp), with all *Mus musculus* genes as a reference list.

## ARRIVE GUIDELINES

The ARRIVE guidelines (https://arriveguidelines.org/) are a checklist of recommendations to improve the reporting of research involving animals. Key elements of the study design should be included below to better enable readers to scrutinize the research adequately, evaluate its methodological rigor, and reproduce the methods or findings.

### Study Design

Details of the number of animals used, sex, age and groups are included in the manuscript text, figures and supplementary files.

### Sample Size

The sample size for this study was determined based on prior experimental experience and standard practices in the field, ensuring sufficient power to detect biologically meaningful effects while accommodating expected variability.

### Inclusion Criteria

The genotype of the mice determined their inclusion in the study.

### Exclusion Criteria

In this atherosclerosis study, mice were excluded if they exhibited dental abnormalities or had body weights that deviated substantially from the expected range. Samples were also excluded from specific analyses when technical issues occurred during processing or data acquisition. Furthermore, animals that developed severe adverse symptoms or required euthanasia, as determined by veterinary assessment, were excluded in accordance with institutional animal welfare guidelines.

### Randomization

The *Ldlr*^-/-^ male mice were randomly assigned to receive bone marrow transplants from either male *Ehd1*^-/-^ mice or age-matched wild type male littermate controls to minimize potential bias in group assignments. During atherosclerotic development, any animals that met the exclusion criteria after randomization were excluded from the study to maintain the integrity of the experimental groups.

### Blinding

Blinding was implemented where appropriate to reduce potential bias: researchers conducting animal experiments were unaware of genotype and treatment assignments, and investigators remained blinded to sample identities during histological analysis, RNA-seq and scRNA-seq analysis, immunofluorescence and quantification.

## Supplemental figures and figure legends

**Figure S1.**
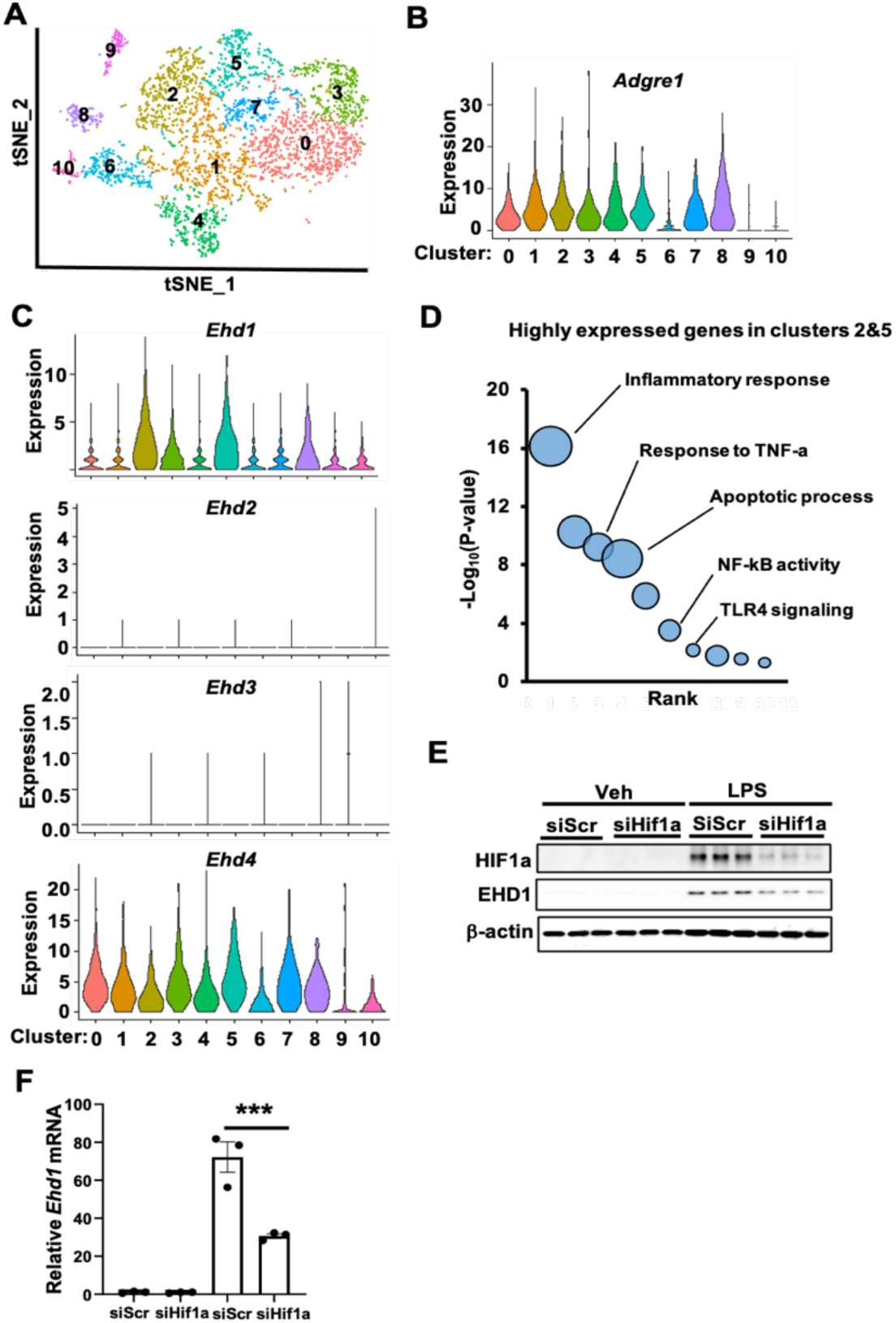
EHD1 is enriched in inflammatory macrophages and regulated by HIF1α. A–D. Reanalysis of published scRNA-seq data of CD45⁺ cells isolated from pooled aortas of *Ldlr^−/−^*mice fed a WD for 12 weeks;^31^ scRNA-seq data was processed with the standard pipeline using Seurat. **A.** T-distributed stochastic neighbor embedding (t-SNE) plot showing the clustering results. **B.** Violin plots showing *Adgre1* expression level in the 11 identified cell clusters. **C.** Violin plots showing *Ehd1*, *Ehd2*, *Ehd3*, and *Ehd4* expression levels in different cell clusters. **D.** GO analysis of the enriched pathways for genes highly expressed in clusters 2 and 5 by comparing the DEGs between clusters 2 and 5 versus the other clusters. **E–F.** Immunoblots of HIF1α and EHD1 (E), and mRNA levels of *Ehd1* (F) in BMDMs transfected with siRNAs, followed by treatment with 100 ng/mL LPS for 24 h (*n* = 3 biological replicates). Data is presented as mean ± SEM. ***p < 0.001 by two-way ANOVA followed by Dunnett’s multiple comparisons test.

**Figure S2.**
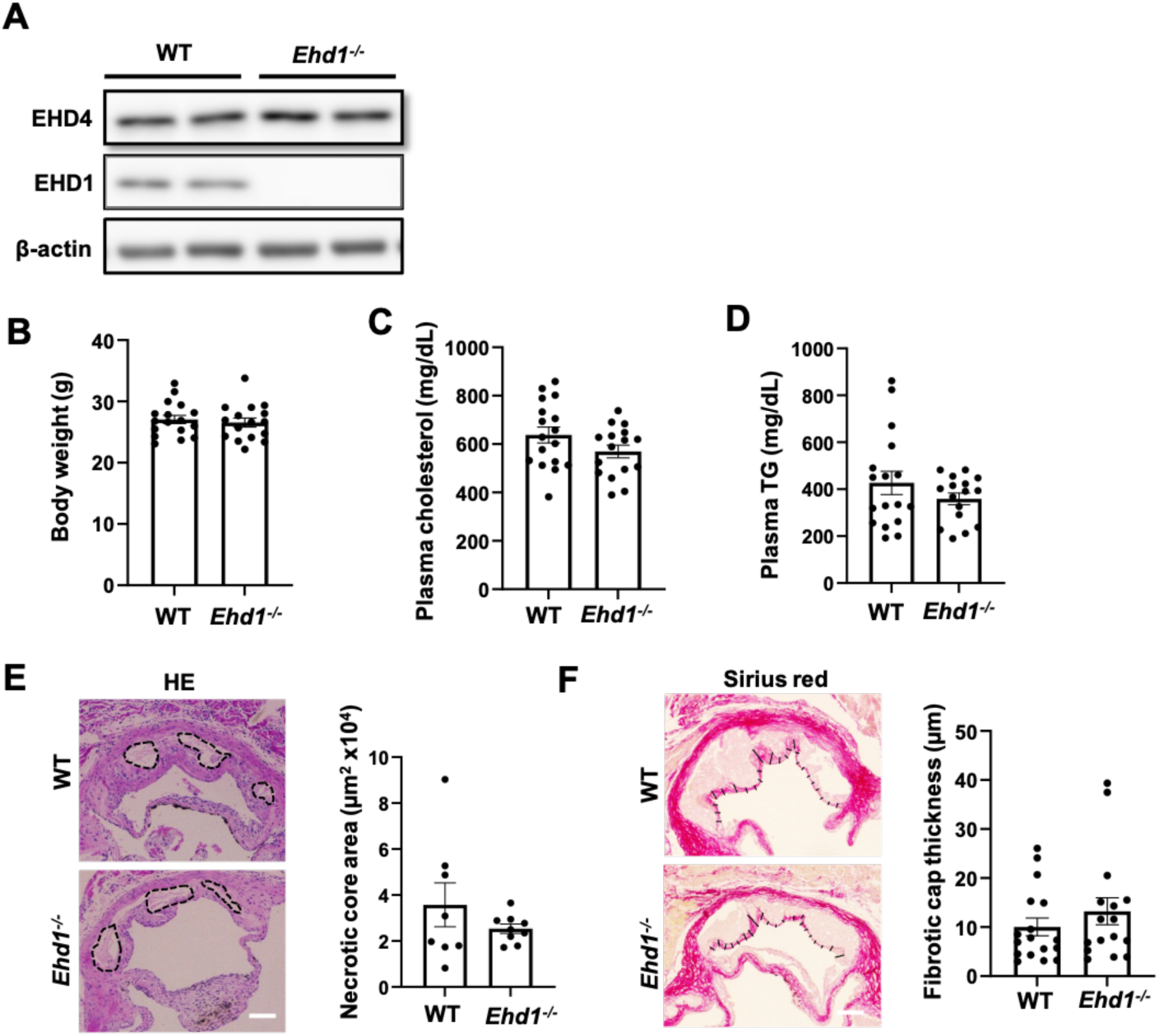
EHD1 deletion has no significant effect on necrotic core area and fibrotic cap thickness in aortic root. **A.** Immunoblots of EHD4 and EHD1 in BMDMs from WT or *Ehd1^-/-^*mice. **B–D.** Body weight (B), plasma cholesterol (C), and triglycerides (D) from WT or Ehd1-/- bone marrow–transplanted male *Ldlr^−/−^*recipients after 12 weeks of WD. **E.** Representative images of necrotic cores in aortic root lesions and quantification with ImageJ (*n* = 16–17/group). Scale bar, 200 μm. **F.** Representative images of fibrotic cap in aortic root lesions, with corresponding quantification. Scale bar, 200 μm. Data is presented as mean ± SEM. n.s., not significant.

**Figure S3.**
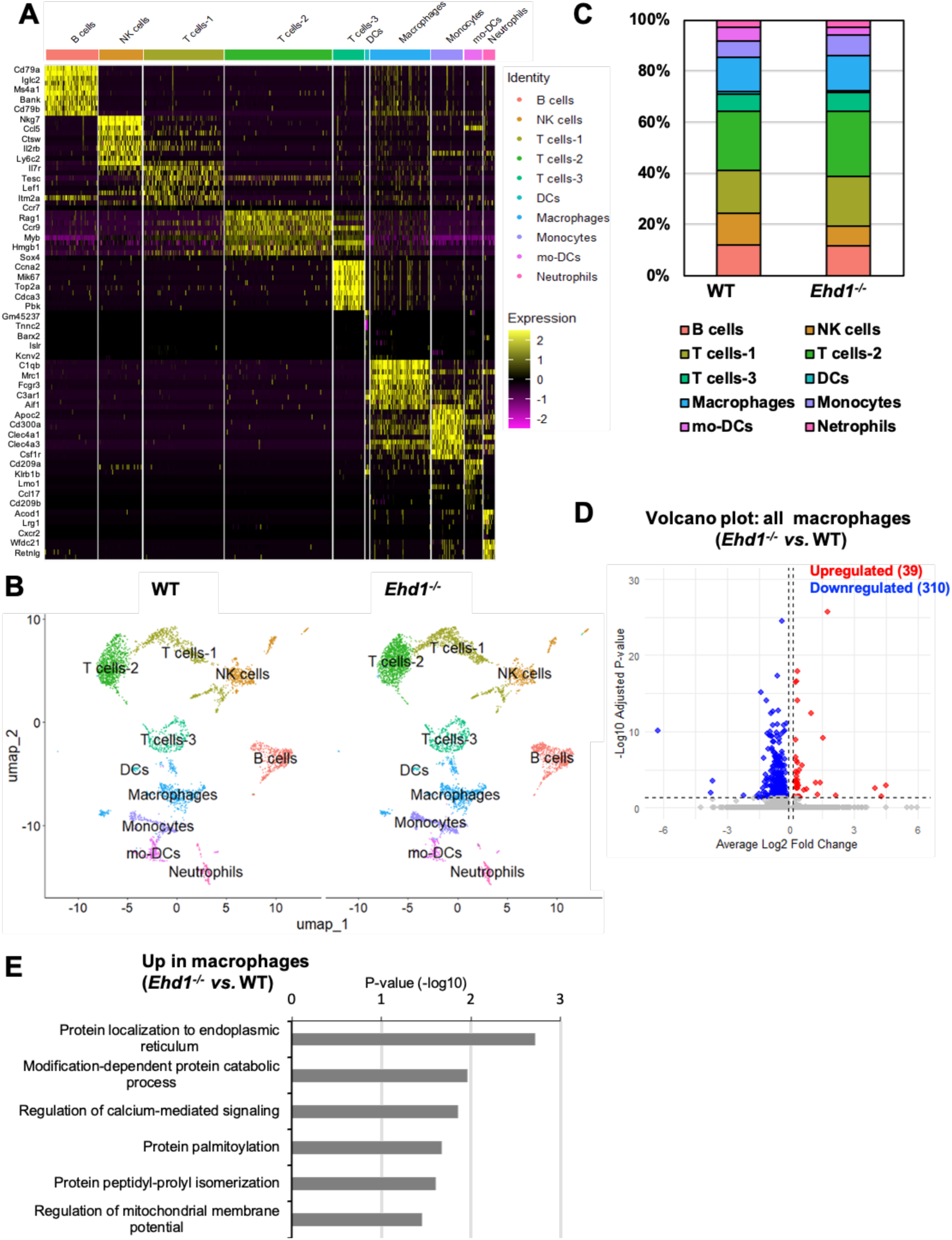
ScRNA-seq analysis of aortic immune cells. **A.** Heatmap showing the top 10 highly expressed genes in each cluster. Five genes were selected from each cluster (shown on the left side). Colors represent normalized gene expression. **B.** UMAP visualization of immune cells separated by the WT and *Ehd1^-/-^* aorta. **C.** Percentage of each immune cell type among total aortic CD45^+^ cells. **D.** Volcano plot showing upregulated and downregulated DEGs in total macrophages isolated from WT and *Ehd1^-/-^* aorta. **E.** GO analysis for the upregulated DEGs in *Ehd1^-/-^ vs.* WT aortic macrophages.

**Figure S4.**
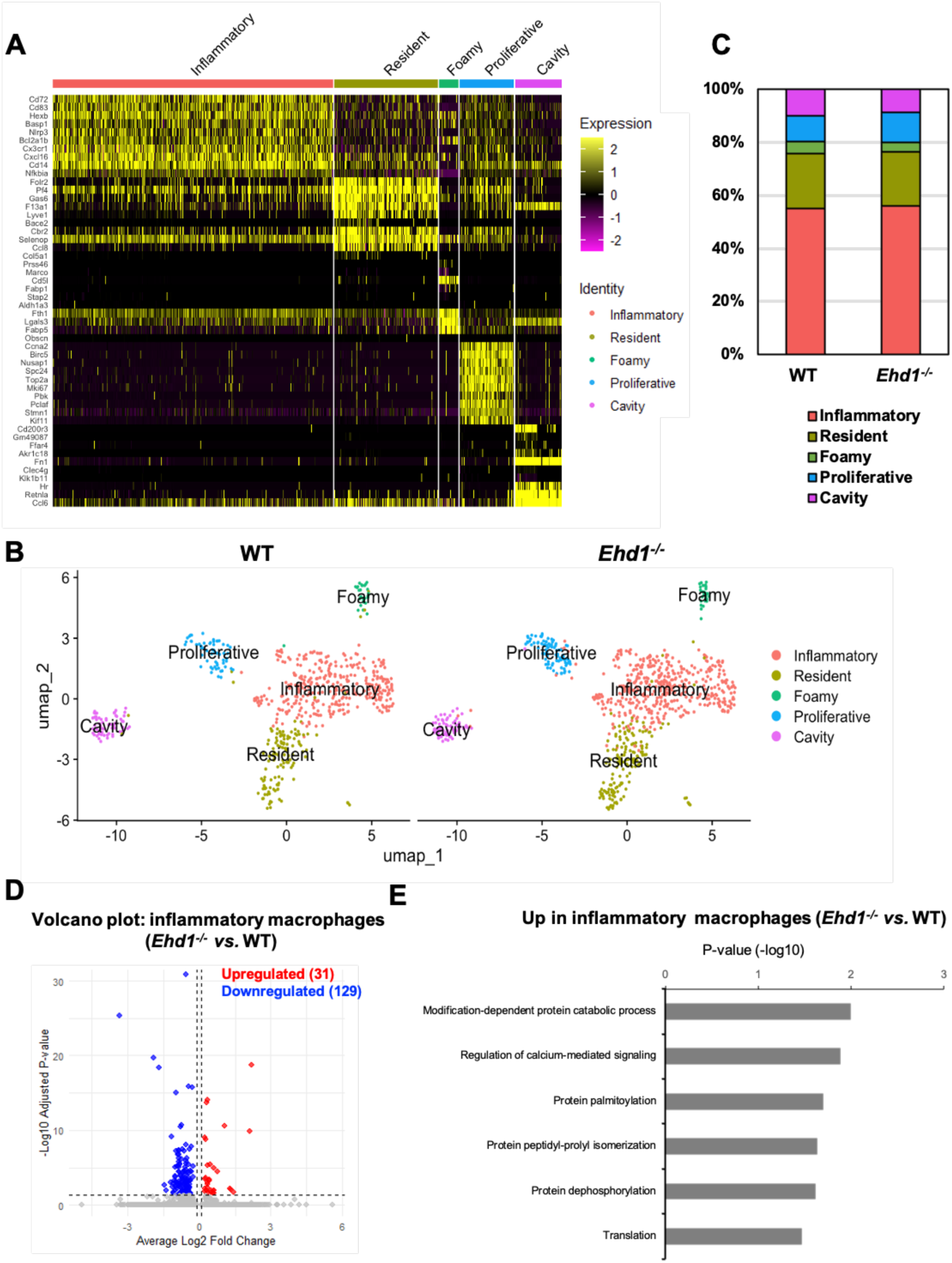
ScRNA-seq analysis of aortic macrophages. **A.** Heatmap showing the top 10 highly expressed genes (on the left side) in each cluster. Colors represent normalized gene expression. **B.** UMAP visualization of macrophage subtypes separated by WT and *Ehd1^-/-^*aorta. **C.** Percentage of each macrophage subset among total aortic macrophages. **D.** Volcano plot showing upregulated and downregulated DEGs in inflammatory macrophages isolated from WT and *Ehd1^-/-^* aorta. **E.** GO analysis for the upregulated DEGs in *Ehd1^-/-^ vs.* WT inflammatory macrophages.

**Figure S5.**
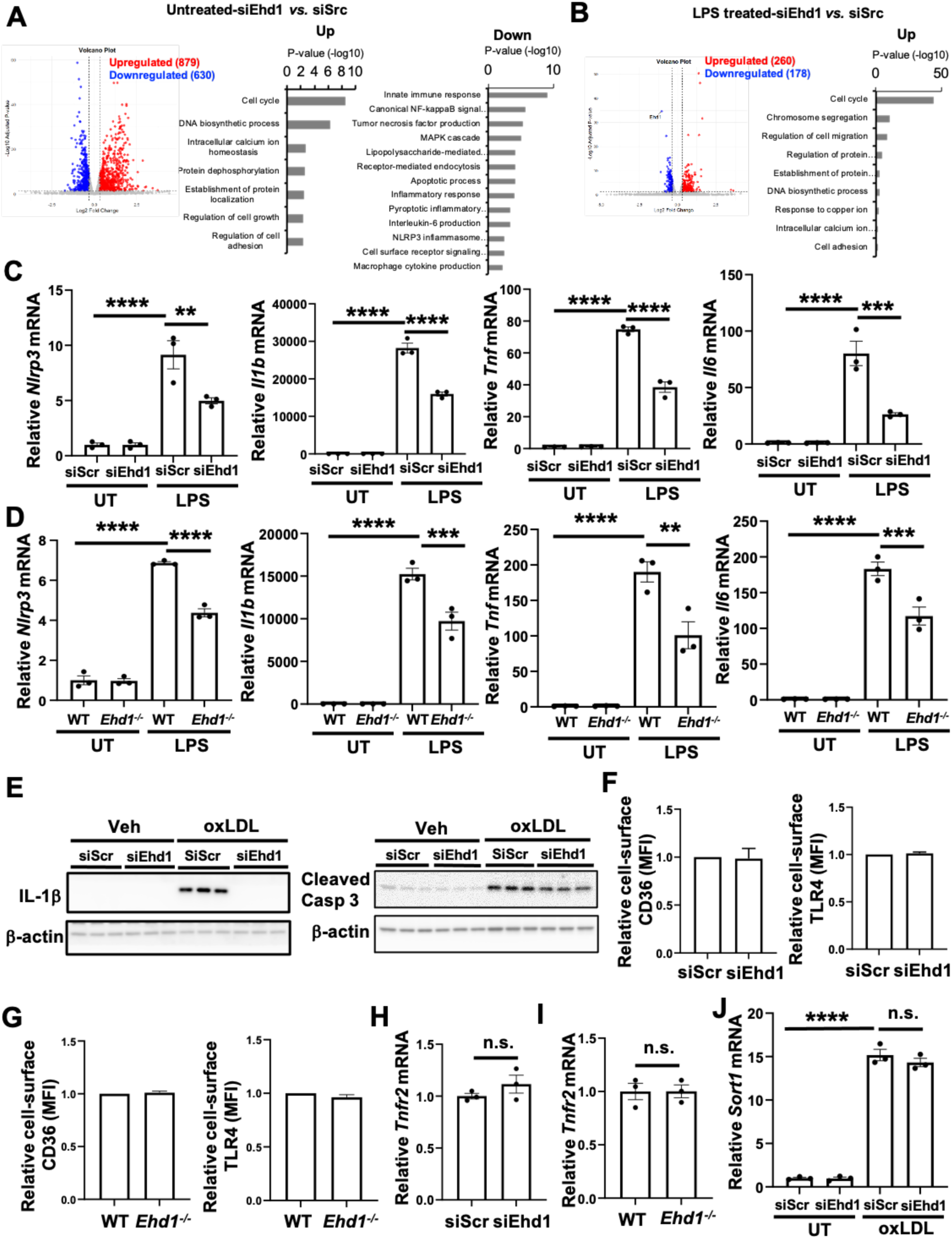
EHD1 deficiency reduces inflammation. **A.** Volcano plot showing upregulated and downregulated DEGs in siEhd1- *vs.* siSrc-treated BMDMs (left). GO analysis for the upregulated and downregulated DEGs in siEhd1- *vs.* siSrc-treated cells (right). **B**. Volcano plot showing upregulated and downregulated DEGs in siEhd1- *vs.* siSrc-treated BMDMs upon LPS treatment (left). GO analysis for the upregulated DEGs in siEhd1- *vs.* siSrc-treated cells, followed by LPS treatment (right). **C–D.** mRNA levels of *Nlrp3*, *Il1b*, *Tnf*, and *Il6* in siEhd1-treated (C) and *Ehd1^-/-^* (D) BMDMs upon LPS treatment (*n* = 3 biological replicates). **E.** Immunoblots of IL1β (left) and Cleaved casp 3 (right) in siRNA-treated BMDMs upon oxLDL treatment. **F–G.** Cell-surface CD36 and TLR4 in siEhd1-treated (F) and *Ehd1^-/-^* (G) BMDMs were detected by flow cytometry (*n* = 3 biological replicates). **H–I.** mRNA levels of *Tnfr2* in siEhd1-treated (H) and *Ehd1^-/-^* (I) BMDMs (*n* = 3 biological replicates). **J.** *Sort1* mRNA levels in BMDMs upon oxLDL treatment (*n* = 3 biological replicates). Data is presented as mean ± SEM. Data was analyzed using one-way ANOVA followed by Dunnett’s multiple comparisons test in C and D, and using Student’s t-test in F, G, H, and I. **p < 0.01, ***p < 0.001, ****p < 0.001. n.s., not significant.

**Figure S6.**
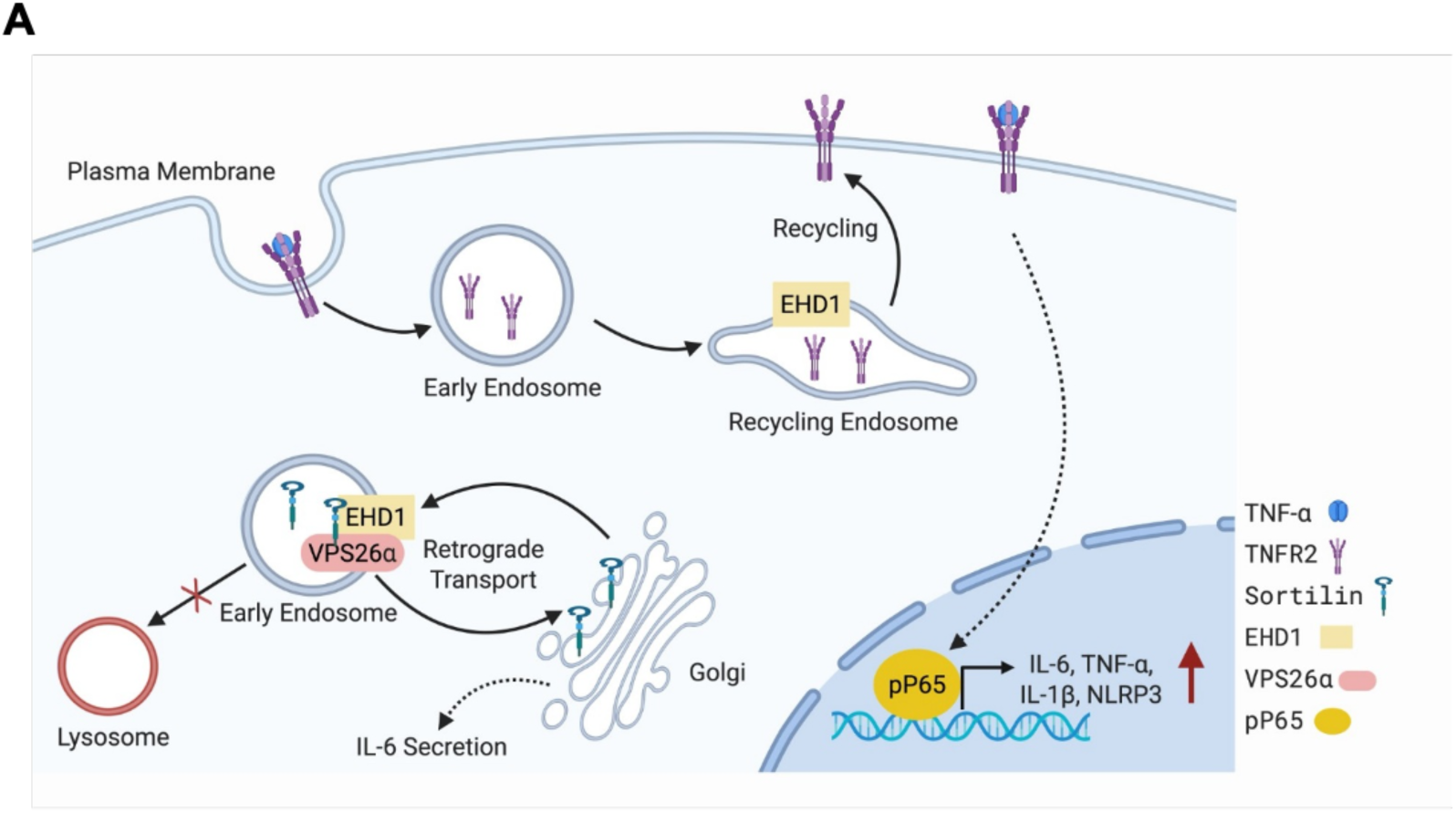
Scheme of the role of EHD1 in regulating inflammation in macrophages. **A.** EHD1 promotes the recycling of endocytosed TNFR2 to the cell surface. Upon engaging with TNF-α, cell-surface TNFR2 signals and activates NF-kB, leading to increased expression of pro-inflammatory mediators, including IL-6, TNF-α, IL-1β, and NLRP3. In addition, EHD1 interacts with retromer, which facilitates the retrograde transport of sortilin from endosomes to TGN and helps stabilize sortilin, a risk factor for atherosclerosis through its role in promoting cytokine secretion.

**Supplemental Table 1.**
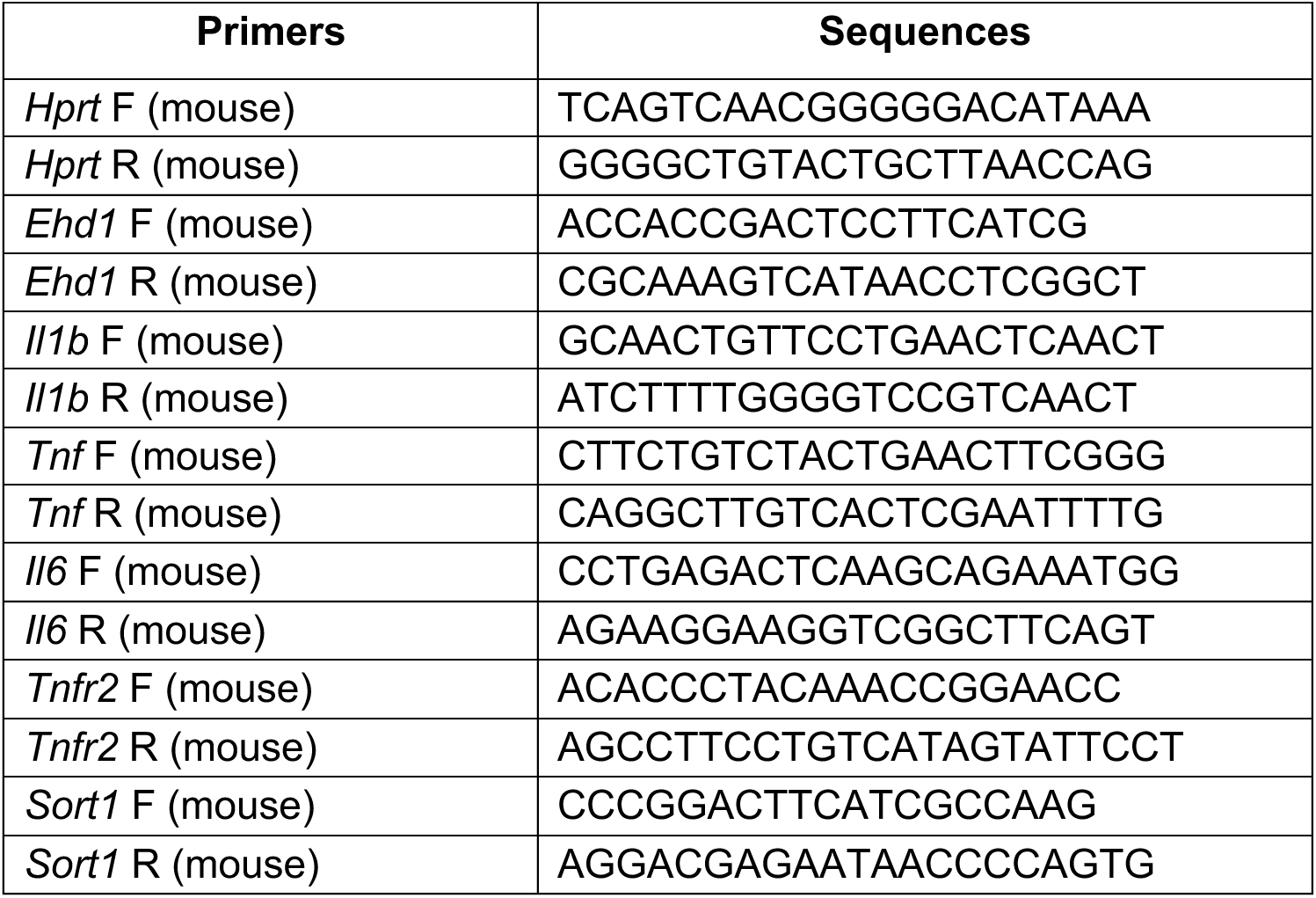
Primers for quantitative RT-PCR.

## Major Resources Table

In order to allow validation and replication of experiments, all essential research materials listed in the Methods should be included in the Major Resources Table below. Authors are encouraged to use public repositories for protocols, data, code, and other materials and provide persistent identifiers and/or links to repositories when available. Authors may add or delete rows as needed.

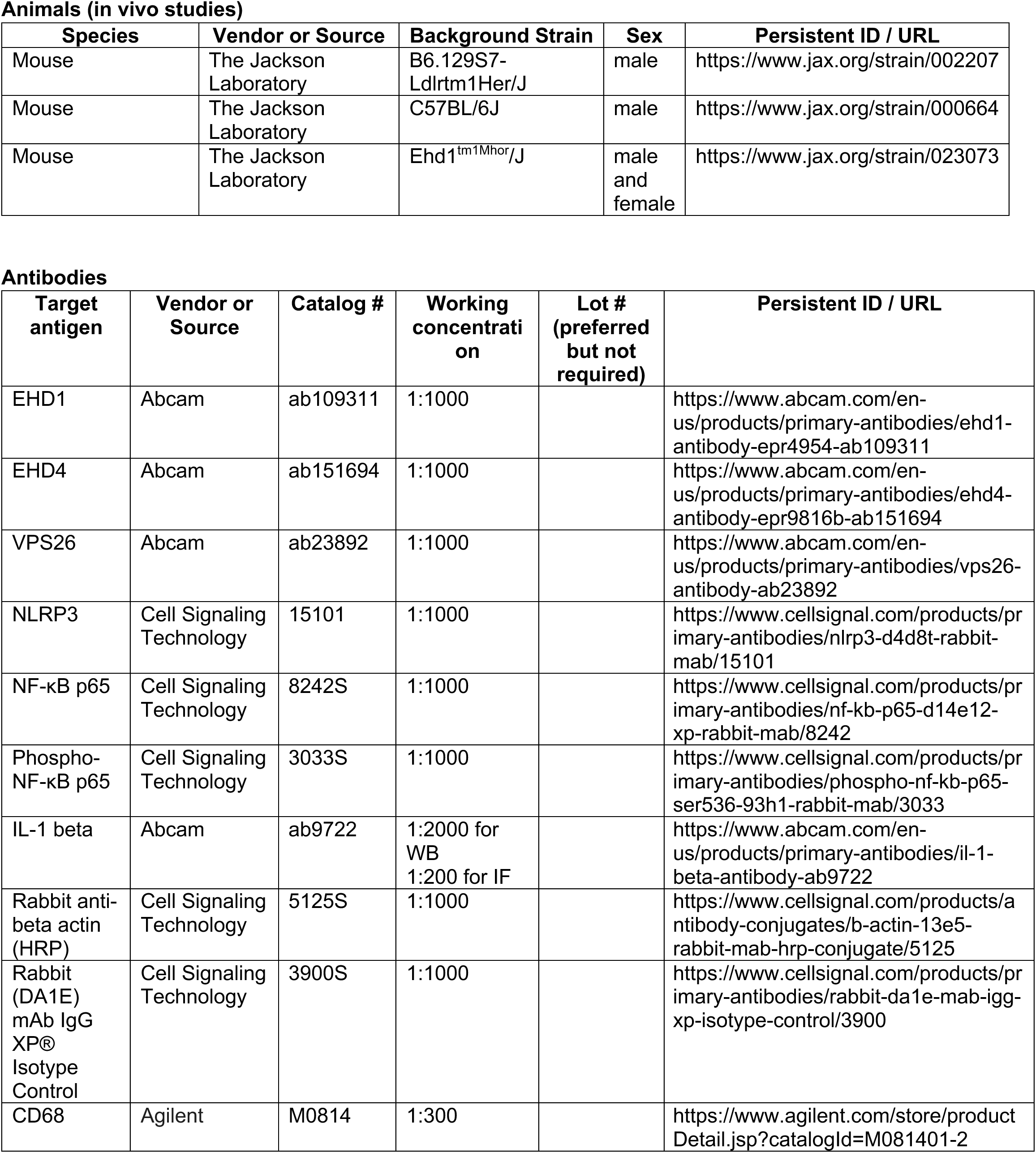
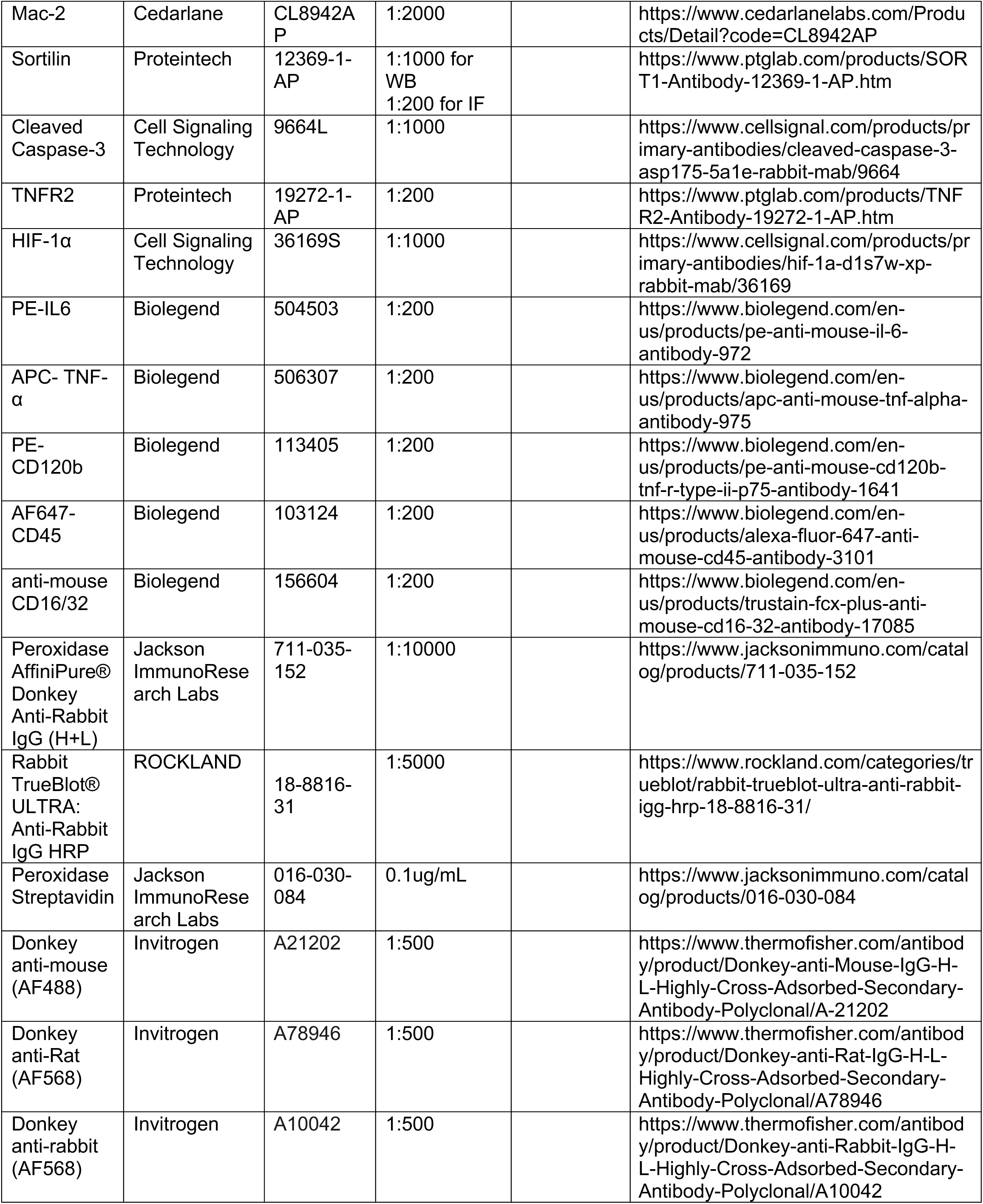
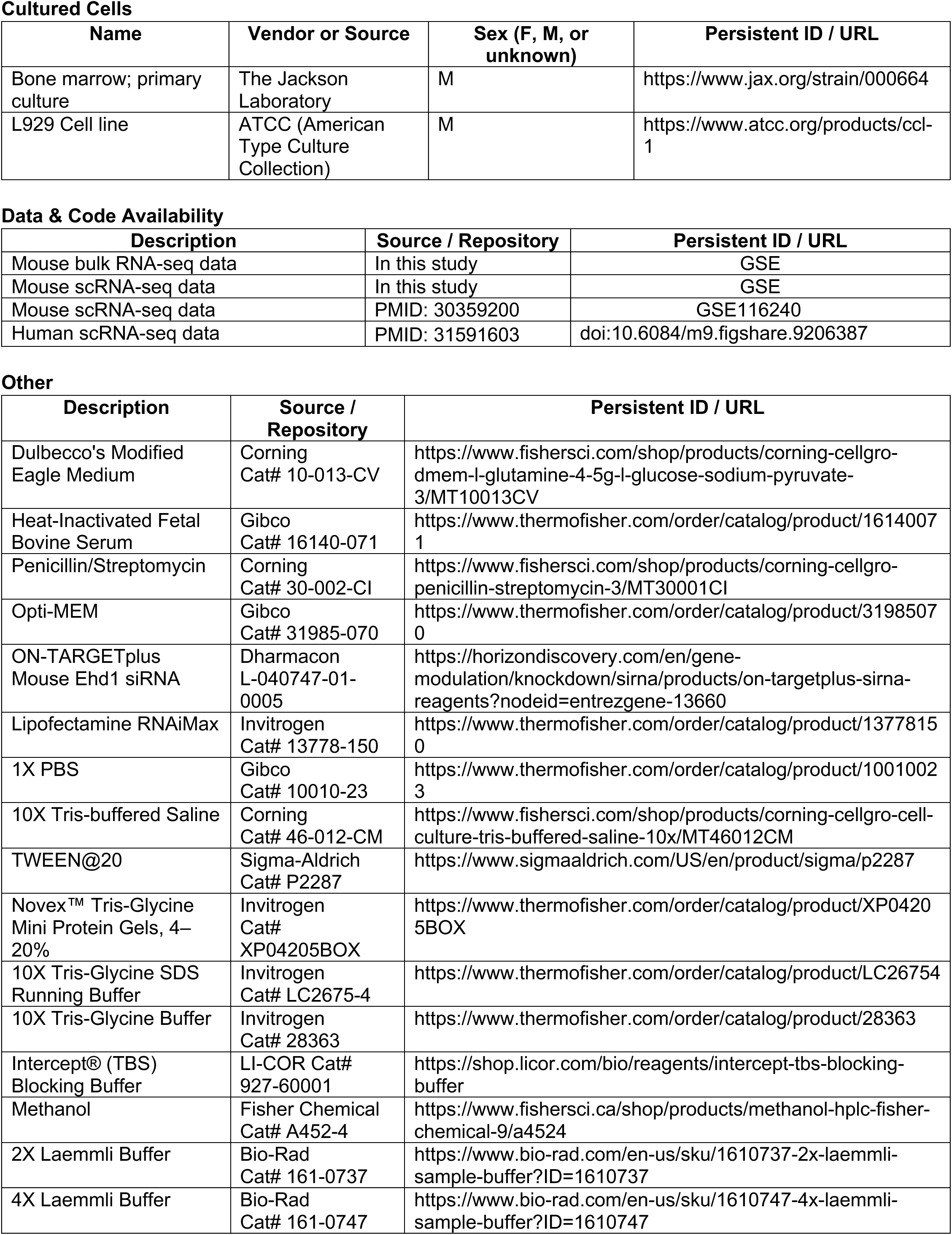
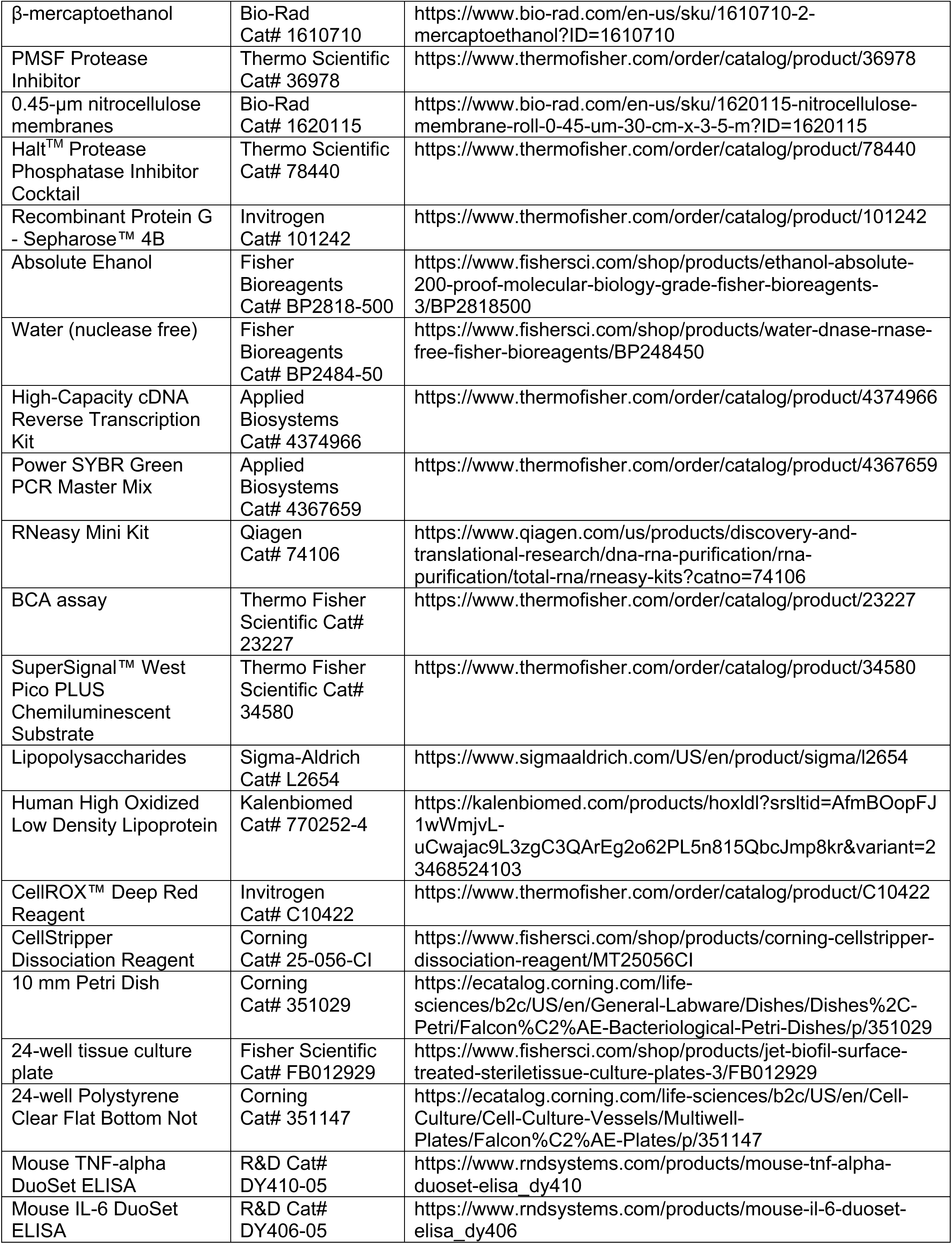
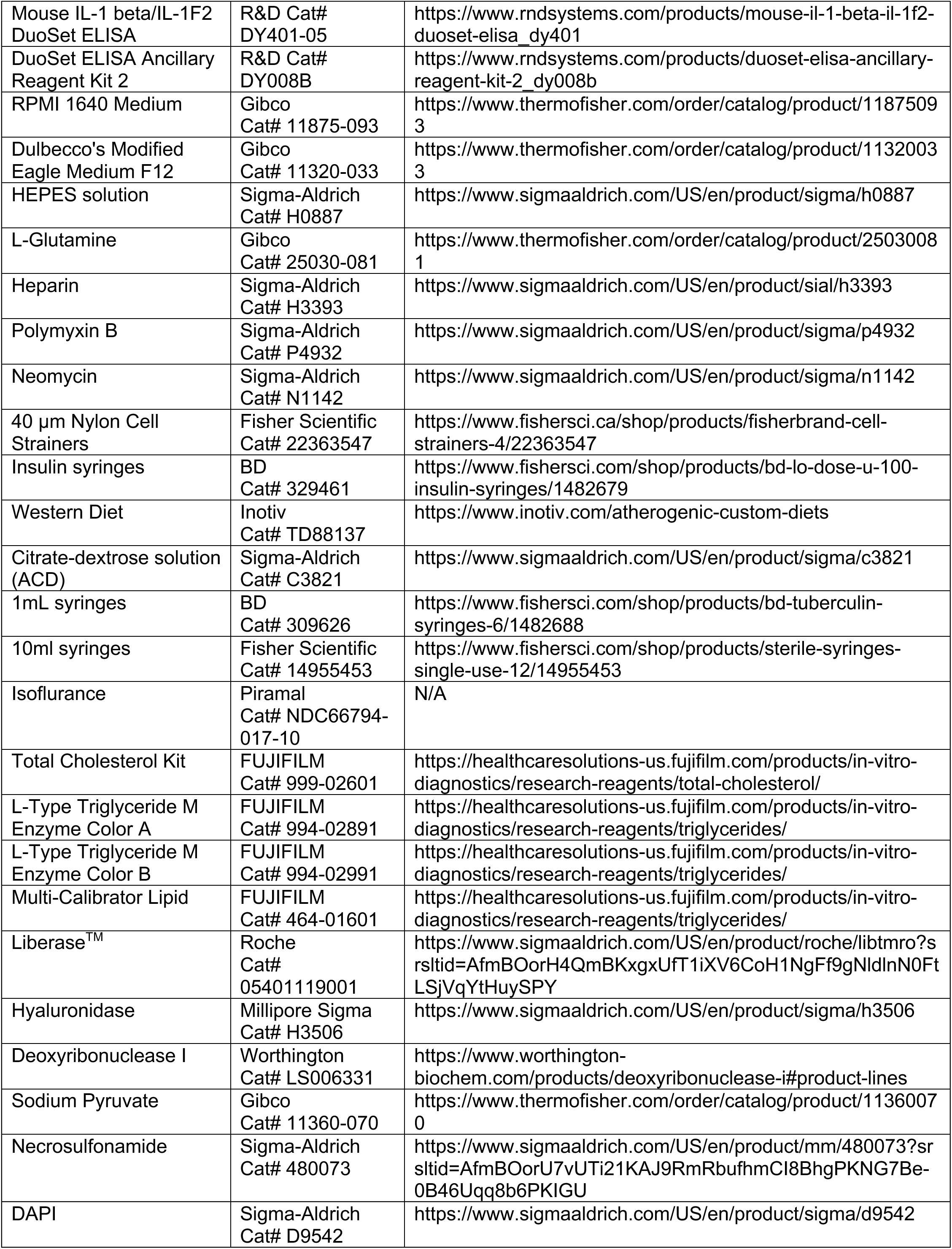
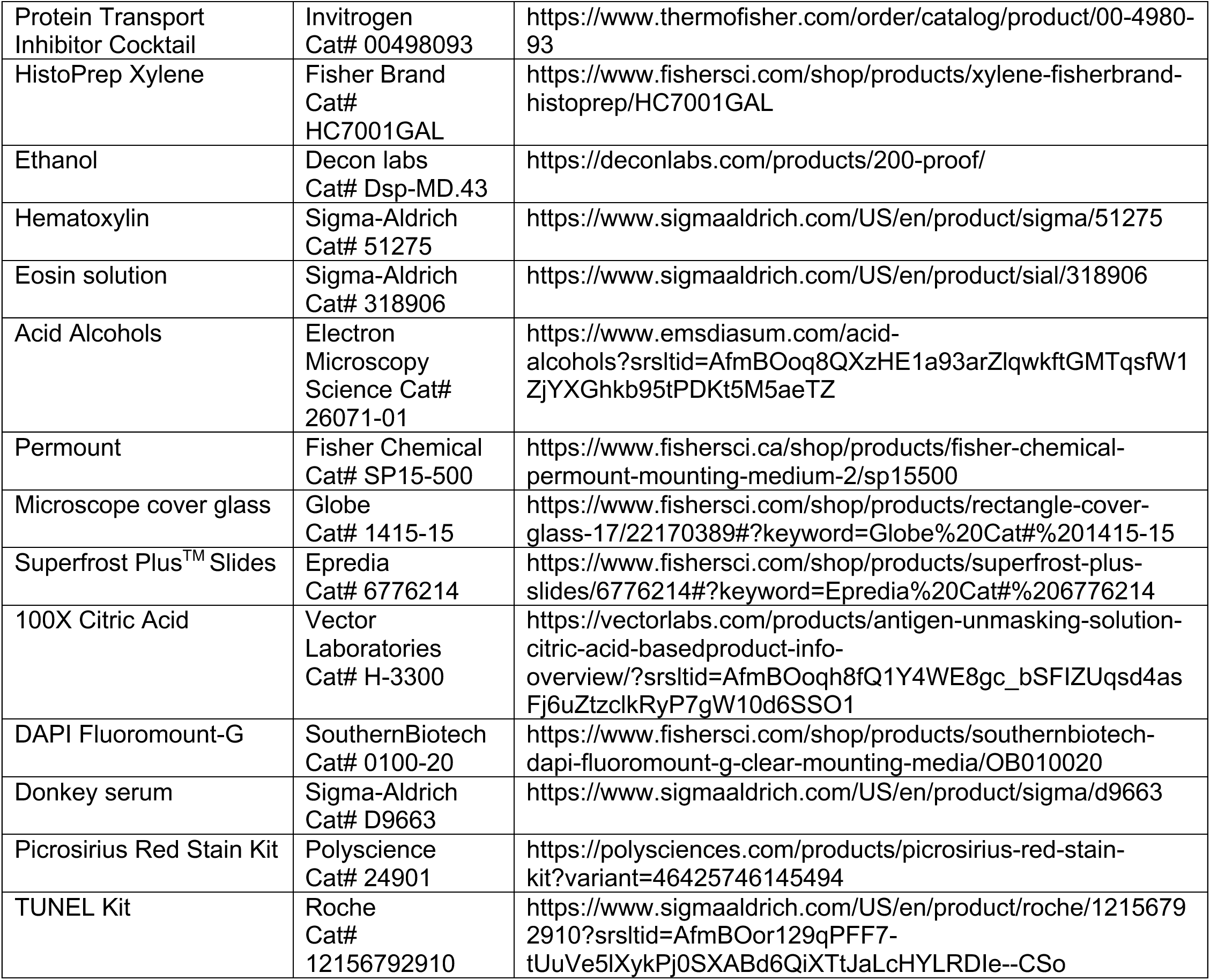

